# Primary Cilia are WNT Transducing Organelles whose Biogenesis is Regulated by a WNT┫ PP1 axis

**DOI:** 10.1101/2022.12.09.519813

**Authors:** Kaiqing Zhang, Fabio Da Silva, Carina Seidl, Michaela Wilsch-Bräuninger, Jessica Herbst, Wieland B. Huttner, Christof Niehrs

**Author notes:** Corresponding author: Christof Niehrs. Co-first authors (Kaiqing Zhang, Fabio Da Silva).

## Abstract

WNT signalling is of paramount importance in development, stem cell maintenance, and disease. WNT ligands typically signal via receptor activation at the plasma membrane to induce β-catenin-dependent gene activation. Here we show that in primary cilia, WNT receptors relay a WNT/GSK3 signal that β-catenin-independently promotes ciliogenesis. Innovations supporting this conclusion are monitoring acute WNT co-receptor activation (phospho-LRP6) and identifying and mutating the LRP6 ciliary targeting sequence. Ciliary WNT signalling inhibits protein phosphatase 1 (PP1) activity, a negative regulator of ciliogenesis, by decommissioning GSK3-mediated phosphorylation of the PP1 regulatory inhibitor subunit PPP1R2. Accordingly, deficiency of WNT/GSK3 signalling by depletion of cyclin Y and cyclin-Y-like protein 1 induces widespread primary cilia defects in mouse embryonic neuronal precursors, kidney proximal tubules, and adult mice preadipocytes. We conclude that primary cilia are WNT ┫ PP1 signalling organelles.

**Graphical Abstract:** **A Localized WNT ┫ PP1 Signalling Axis Promotes Ciliogenesis** The WNT co-receptor LRP6 localizes to the ciliary membrane, where it is phospho-primed via a CCNY/L1-dependent CDK (not shown). WNT signalling inhibits GSK3 (not shown) and leads to inhibition of Protein phosphatase 1, a negative regulator of ciliogenesis. Right, CCNY/L1 deficiency disrupts the WNT ┫ PP1 signalling axis, leading to ciliary defects.

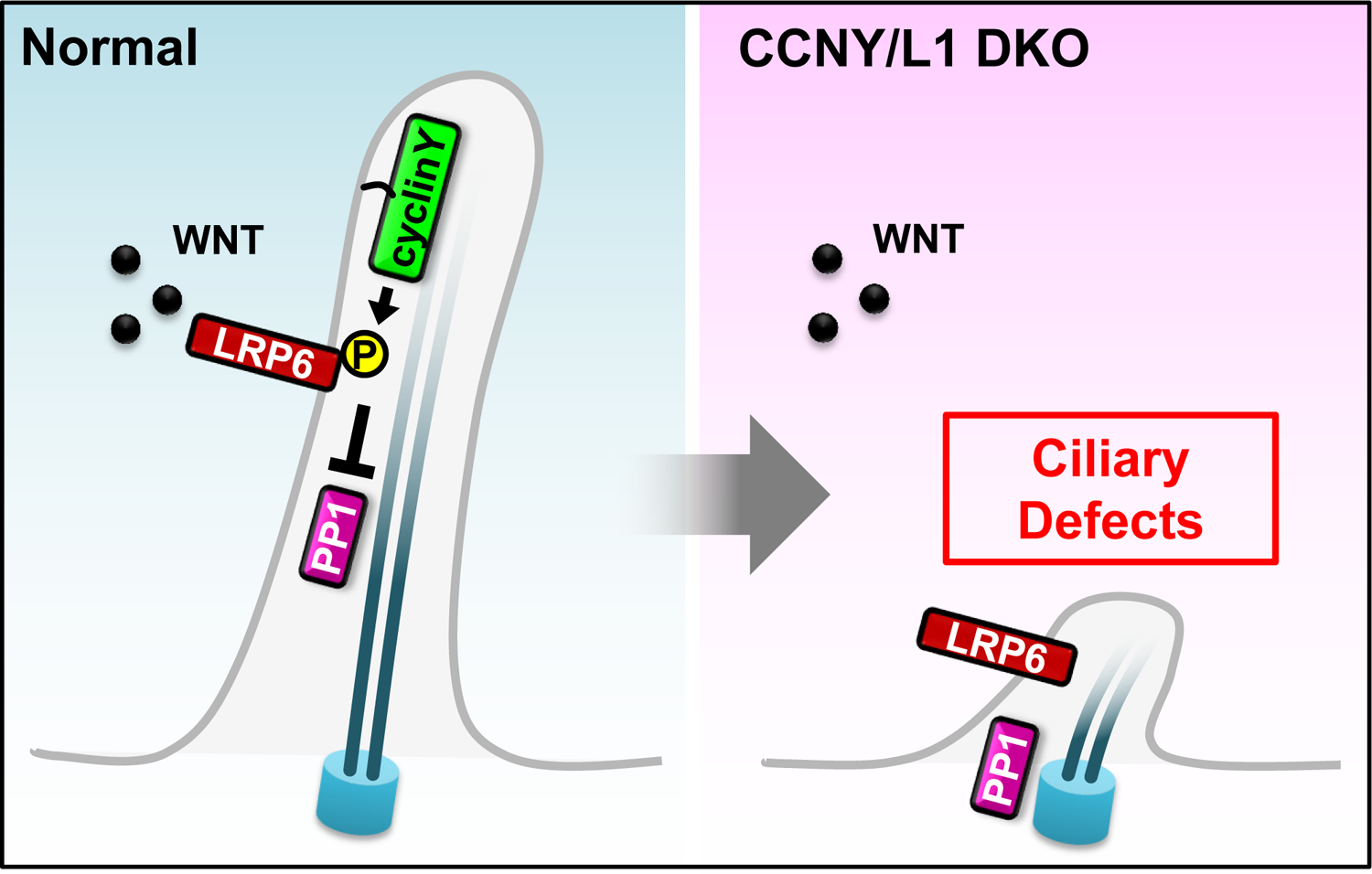

## INTRODUCTION

Does a growth factor trigger the same response through the plasma membrane as it does through cilia? In the case of Hedgehog signalling, emerging evidence indicates that cilia can transduce a response that is distinct from that of the plasma membrane (Truong et al., 2021). Here we investigate the possibility of distinct ciliary signalling for WNT, a growth factor paramount in development and disease. Canonical WNT/β-catenin-dependent signalling proceeds via classical transmembrane signalling across the plasma membrane. Key WNT receptors and co-receptors include members of the Frizzled (FZD) and low-density lipoprotein receptor-related protein (LRP5 and 6) classes (Niehrs, 2012; Jeong et al., 2021). LRP6 and FZD transmembrane receptors bind WNT ligands to induce a cytoplasmic signalling cascade that stabilizes the transcriptional coactivator β-catenin through inhibition of a destruction complex containing glycogen synthase kinase 3 (GSK3) (Kikuchi et al., 2011; De Herreros and Duñach, 2019). GSK3 inhibition leads to dephosphorylation of cytoplasmic β-catenin, releasing the protein from proteasomal degradation, allowing it to enter the cell nucleus, and engage a transcription complex to regulate downstream target genes (Nusse and Clevers, 2017). WNT signalling across the plasma membrane via FZD and LRP6 is well-documented (MacDonald and He, 2012), including by live cell imaging (Bilic et al., 2007; Holzer et al., 2012).

Continuous with-but insulated from the plasma membrane are cilia, organelles that are chemosensory and which transduce certain growth factor signals. Ciliogenesis, the growth and development of cilia, depends on a dynamic equilibirum of ciliary assembly and disassembly (Bernabé-Rubio and Alonso, 2017; Malicki and Johnson, 2017). There are three classes of cilia, motile and non-motile (primary) cilia as well as flagella, and dysfunction of all three cilia classes leads to a spectrum of diseases known as ciliopathies (Valente et al., 2014; Braun and Hildebrandt, 2017; Reiter and Leroux, 2017). The coordinated movement of motile cilia moves fluids across epithelia or, in the case of flagella, propels cells forward. Primary cilia have a sensory function and their membranes harbor numerous receptors that sense fluid flow and content. They concentrate signalling receptors and are prominent signalling hubs for major growth factor families (Hilgendorf et al., 2016; Anvarian et al., 2019; Nachury and Mick, 2019), including PDGF, EGFR, IGF, Notch, and Hh (Goetz and Anderson, 2010).

In regards to the canonical WNT pathway, there exists a three-fold relationship between cilia and WNT/β-catenin signalling. First, it is well established that WNT/β-catenin signalling promotes cilia formation in zebrafish, frog, and mouse via transcriptional gene activation of ciliogenesis regulators such as of the transcription factor *Foxj1* (Brechbuhl et al., 2011; Caron et al., 2012; Walentek et al., 2012; Li Song et al., 2014; Walentek et al., 2015; Walentek et al., 2015b; Kyun et al., 2020; Zhang et al., 2020). Second, a body of work manipulating ciliary components and monitoring the effects on WNT signalling supports an inhibitory function of primary cilia on Wnt/β-catenin signalling (Simons et al., 2005; Gerdes et al., 2007; Corbit et al., 2008; McDermott et al., 2010; Wiens et al., 2010; Lancaster et al., 2011; Liu et al., 2014; Zingg et al., 2018), although the question whether cilia restrain canonical WNT signalling is controversial as deficiency of multiple ciliary core components in e.g. zebrafish and mutant mice has no effect on WNT signalling (Scholey and Anderson, 2006; Huang and Schier 2009; Ocbina et al., 2009). Of note, the contentious inhibitory function of primary cilia components on Wnt/β-catenin signalling is occasionally referred to as ‘ciliary WNT signalling’. The term is misleading as it may imply that cilia respond to- and transduce WNT signalling, while the converse is meant, cilia inhibit WNT signalling. Third, a number of signal transducing proteins acting in the WNT pathway also locate to the cilium, including Dishevelled (DVL), FZD, and GSK3 (Thoma et al., 2007; Veland et al., 2013; Stypulkowski et al., 2021) but whether their presence reflects true ‘ciliary WNT signalling’ remains speculative. The reason is that approaching ciliary WNT signalling requires either monitoring a direct ciliary response to WNT ligands or experimentally uncoupling ciliary-from plasma membrane signalling, neither of which is documented. Consequently, despite a substantial body of work the key question remains open: Is there WNT signalling within cilia? Given the widely held notion that the main output of WNT signalling is a change in β-catenin-mediated transcription, what may be the cellular function of such intra-ciliary WNT signalling, and what physiological significance could such intra-ciliary WNT signalling have *in vivo*?

A clue came from our previous report, which showed that WNT triggers a non-transcriptional WNT/GSK3 response in mouse sperm flagella that is essential for sperm maturation (Koch et al., 2015). First, this observation supports the emerging idea that the canonical WNT signalling cascade induces a much richer response than just β-catenin activation. This is because GSK3 inhibition by WNT disables phosphorylation of hundreds of other proteins besides β-catenin (Taelman et al., 2010; Acebron et al., 2014), and thereby directly affects a wide array of cellular functions besides transcription (Inoki et al., 2006; Huang et al., 2015; Kim et al., 2015; Demagny and De Robertis, 2016; Madan et al., 2018; Hinze et al., 2019; Tejeda-Munoz et al., 2019; Lin et al., 2020; Da Silva et al., 2021). Second, the link between flagella and WNT signalling raised the intriguing possibility that WNT/GSK3 signalling may not be limited to highly specialized spermatocytes, and may also operate in primary cilia of various cell types.

To address the possibility of a role of WNT/GSK3 in primary cilia, we leveraged the specificity of cyclin Y (CCNY) and its close homolog cyclin Y-like 1 (CCNYL1; collectively CCNY/L1) for the β-catenin-independent WNT/GSK3 pathway. CCNY/L1 are mitotic cyclins that activate the cyclin-dependent kinase family members CDK14 and CDK16, which phosphorylate and prime the WNT co-receptor LRP6 for incoming WNT ligands (Davidson et al., 2009; Acebron et al., 2014). Depleting CCNY/L1 offers the opportunity to inhibit and study WNT/GSK3 signalling on the background of largely intact β-catenin function (Acebron et al., 2014; Huang et al., 2015; Koch et al., 2015; Da Silva et al., 2021). Important for this study, CCNY/L1 mediate WNT/GSK3 signalling not only in dividing cells but also in non-dividing sperm to induce flagella-maturation (Koch et al., 2015), consistent with non-cell cycle functions of the CDK Network (Liu et al., 2019).

Here we show that *Ccny/l1* deficiency leads to primary cilia defects in mouse embryos, akin to neurological ciliopathies. In adult mice, we provide a direct link between WNT/GSK3 signalling, primary cilia, and fat homeostasis. We show that primary cilia are WNT transducing organelles harbouring cilia-localized LRP6 co-receptors that respond to- and transmit WNT signals. Live-cell imaging reveals ciliary localization of LRP6 from the very onset of ciliogenesis. Notably, we identify a ciliary targeting signal in LRP6 that it is essential for ciliogenesis, thereby for the first time functionally dissociating ciliary-from cytoplasmic WNT signalling.

We unravel a ciliary signalling cascade where WNT/GSK3 signalling independently of β-catenin promotes ciliogenesis by inhibiting protein phosphatase 1 (PP1), a direct ciliary regulator. In conclusion, we propose that primary cilia are WNT ┫ PP1 signalling organelles. Our insights how WNT signalling directly promotes ciliogenesis may aid in understanding the etiology of ciliopathies.

## RESULTS

### Exencephaly and Ciliogenesis Defects in *Ccny/l1^-/-^* Mouse Embryos

To investigate the role of WNT/GSK3 signalling in primary cilia in mouse, we analyzed mice deficient for cyclin Y and cyclin Y-like 1 (collectively *Ccny/l1*), key regulators of the pathway. *Ccny/l1* double knockout (DKO) embryos displayed premature lethality at E14.5 (Da Silva et al., 2021). To avoid that the results of our analyses might reflect non-specific effects resulting from early lethality, we analysed DKO and littermate control embryos at E13.5. Interestingly, 16% of DKO embryos (n=60) displayed cranial exencephaly (Figure 1A), a characteristic malformation in humans with primary ciliopathies (Valente et al., 2014). Exencephaly was observed at similar frequencies during earlier time-points (E11.5 (20%; n=10) and E12.5 (20%; n=10)), suggesting that it results from impaired neural tube closure, which is completed around E9.5 in mice (Juriloff and Harris, 2000).

**Figure 1.**
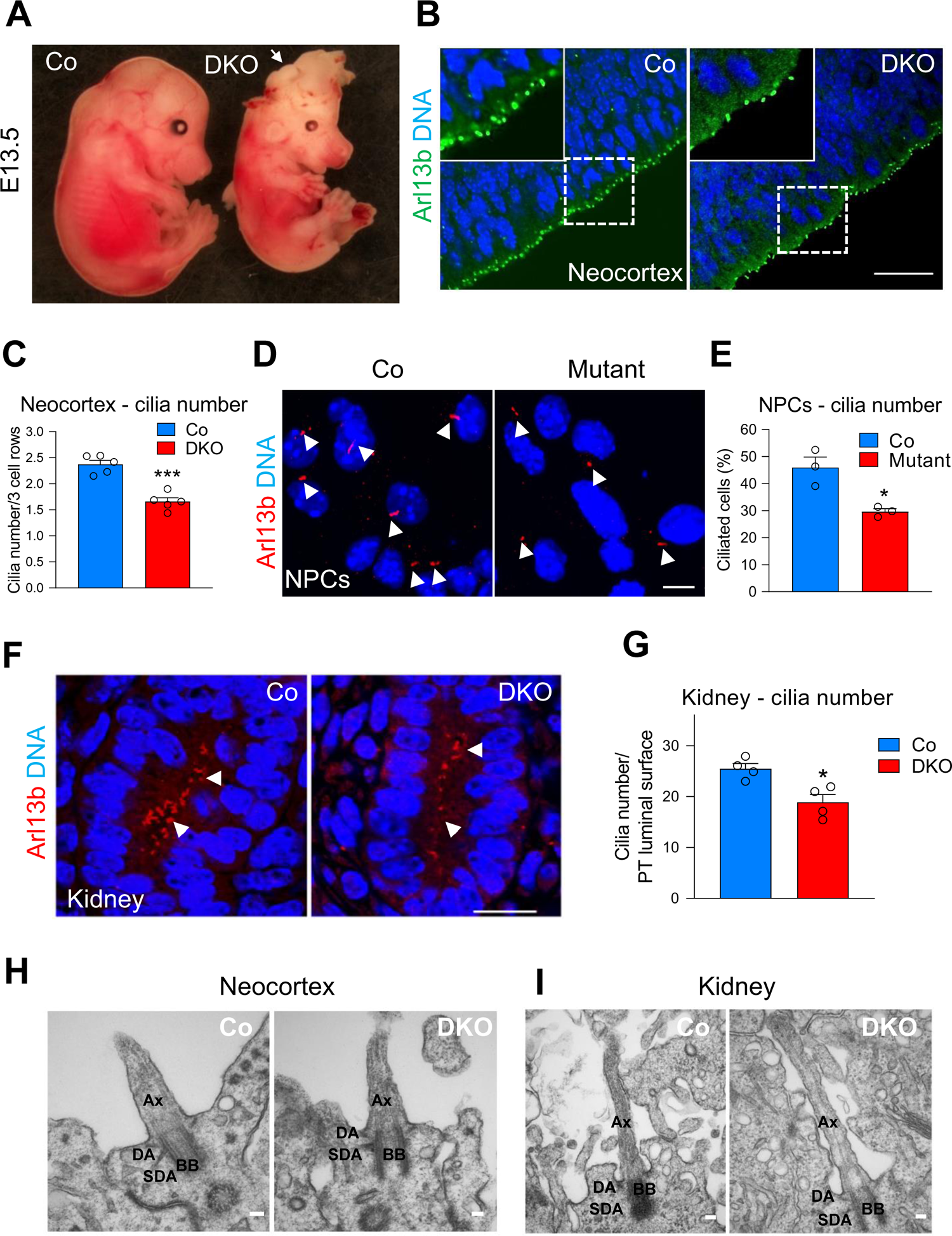
Exencephaly and Ciliogenesis Defects in *Ccny/l1^-/-^* Mouse Embryos. **(A)** *Ccny/l1* double knockout (DKO) E13.5 mouse embryos exhibit cranial exencephaly (white arrow) (n=60 embryos analyzed). **(B)** Immunofluorescence (IF) for the ciliary membrane protein Arl13b in the neocortex of E13.5 embryos. White dashed boxes are magnified in insets and depict the ciliated apical membrane of the neocortex. DNA visualized by Hoechst staining. Scale bar 20 µm. **(C)** Quantification of **(B)**. Cilia number was normalized to the first three layers of cells in the neocortex. Quantification was performed on 5 control (Co) and DKO embryos obtained from 3 different litters. Data are means ± SEM. **(D)** Arl13b IF on control (*Ccny*^+/-;^ *Ccnyl1*^-/-^; shRNA *Co*) and mutant (*Ccny*^+/-^; *Ccnyl1*^-/-^; shRNA *Ccny*) monolayer cultures of neural progenitor cells (NPCs) isolated from E13.5 forebrains. Cilia depicted by white arrowheads. Scale bar 5 µm. **(E)** Quantification of **(D)**. Data are means ± SEM (experiment performed twice in triplicates, representative experiment shown). **(F)** Arl13b IF on E13.5 kidney proximal tubules. Arrowheads indicate ciliated apical membrane. Scale bar 10 µm. **(G)** Quantification of **(F)**. Cilia number in developing proximal tubules (PT) normalized to the perimeter of the luminal surface. Data are means ± SEM (n=4 embryos, 4 litters). **(H)** Transmission electron microscopy (TEM) analysis of representative apical cilia in control and DKO neocortex. No obvious differences in morphology were observed. Ax = axoneme, DA = distal appendage, SDA = subdistal appendage, BB = basal body. **(I)** TEM analysis of control and DKO apical cilia in kidney proximal tubules reveals no major differences in morphology. Data information: Unpaired two-tailed t-test for all statistical analyses: ns, not significant; * p < 0.05, *** p < 0.001.

Since neural tube closure defects are associated with defects in primary cilia (Valente et al., 2014), we examined the levels of primary cilia in the neural tube of E13.5 DKO embryos. We stained sections of the neocortex for Arl13b (ADP-ribosylation factor-like protein 13b), which marks the ciliary membrane. DKO embryos displayed a 25% decrease (p = 0.0001) in the number of primary cilia at the apical membrane of the progenitor cells in the ventricular zone compared to controls (Figure 1B-C). We confirmed the results by IF for the ciliary axoneme marker Ift81 (Intraflagellar transport protein 81), which revealed a similar decrease (−28%, p = 0.02) in primary cilia number (Figure S1A-B). Reduced cilia number was also detected in the ventral neural tube of E9.5 embryos, suggesting Ccny/l1 are required for cilia formation throughout embryogenesis (Figure S1C-D). To recapitulate the results *in vitro*, we isolated and cultured neural progenitor cells from E13.5 *Ccnyl1*-/- embryos, transduced them with a lentivirus expressing shRNA against *Ccny*, and performed IF for Arl13b. Consistently, *Ccny/l1-* deficient progenitors developed fewer primary cilia compared to shRNA controls (−36%, p = 0.04) (Figure 1D-E). This result indicates that the cilia defects in DKO embryos reflect an acute, direct requirement for Ccny/l1 in neural progenitors.

Another organ in which primary cilia play an essential role is the kidney, and polycystic kidney disease, a debilitating inherited disorder, is highly associated with mutations in genes required for primary cilia formation (Kathem et al., 2014). To analyze if *Ccny* and *Ccnyl1* are expressed in the developing kidney at E13.5, we carried out IF with previously validated antibodies (Da Silva et al., 2021) and detected enrichment of both proteins in the apical cell cortex or membrane of developing proximal tubules (Figure S1E). Ccny proximal tubule staining was absent in DKO embryos, confirming antibody specificity (Figure S1F). IF for Arl13b in DKO proximal tubules showed reduced primary cilia number (−26%, p=0.02) when compared to controls (Figure 1F-G). The effects of Ccny/l1 on ciliogenesis in the embryonic neocortex and kidney were related to cilia number but not their integrity, as no obvious defects in cilia morphology or length were detected by transmission electron microscopy (TEM) of DKO neocortex and kidneys (Figure 1H-I), and by scanning electron microscopy (SEM) analysis of DKO forebrains (Figure S1G). Altogether, these results demonstrate that Ccny/l1 promote primary cilia formation in the embryonic mouse brain and kidney and that their deficiency recapitulates a classical ciliopathy, exencephaly.

### Lipodystrophy in *Ccny^-/-^* Mice due to Reduced Ciliogenesis

A previous study reported that single *Ccny^-/-^* mice show dysfunctional lipid metabolism and develop less adipose tissue due to reduced differentiation of preadipocyte progenitors (An et al., 2015). Interestingly, preadipocyte progenitors represent the major ciliated cell type in fat tissue and depletion of cilia in preadipocytes impairs their ability to differentiate *in vitro* and *in vivo* (Zhu et al., 2009; Dalbay et al., 2015; Hilgendorf et al., 2019). Hence, could decreased adipogenesis in *Ccny^-/-^* mice be due to impaired ciliogenesis?

To test this possibility, we first confirmed the adipogenesis defects in *Ccny* mutants (An et al., 2015). As reported, *Ccny^-/-^* mice were smaller (Figure 2A), exhibited decreased epididymal, inguinal, and interscapular fat mass (Figure 2B-C) and displayed smaller adipocytes in epididymal fat pads (Figure 2D). We analyzed primary cilia in preadipocytes, a marker for which is Pdgfrα (Berry and Rodeheffer, 2013). IF analysis of epididymal fat pads for Arl13b^+^ preadipocytes in *Ccny^-/-^* mice revealed a marked decrease in cilia number (−38%, p = 0.01) (Figure 2E-F). Consistently, acute siRNA knockdown of *Ccny* in preadipocyte 3T3-L1 cells decreased cilia number (Figure 2G-H) and length (Figure 1N). In addition, acute siRNA knockdown of *Ccny* in 3T3-L1 preadipocytes followed by induction of differentiation reduced expression of the adipocyte markers and *Fabp4* (fatty acid binding protein 4) and *Cebpa* (CCAAT/enhancer-binding protein alpha) (Figure S1H). In stark contrast, si*β-catenin* enhanced adipogenesis (Figure S1H), consistent with the well-established negative role of WNT/β-catenin signalling on adipogenic differentiation (Ross et al., 2000; Bennett et al., 2002; Christodoulides et al., 2008). These results support that Ccny promotes primary cilia formation in adult preadipocytes and that its deficiency leads to lipodystrophy.

**Figure 2.**
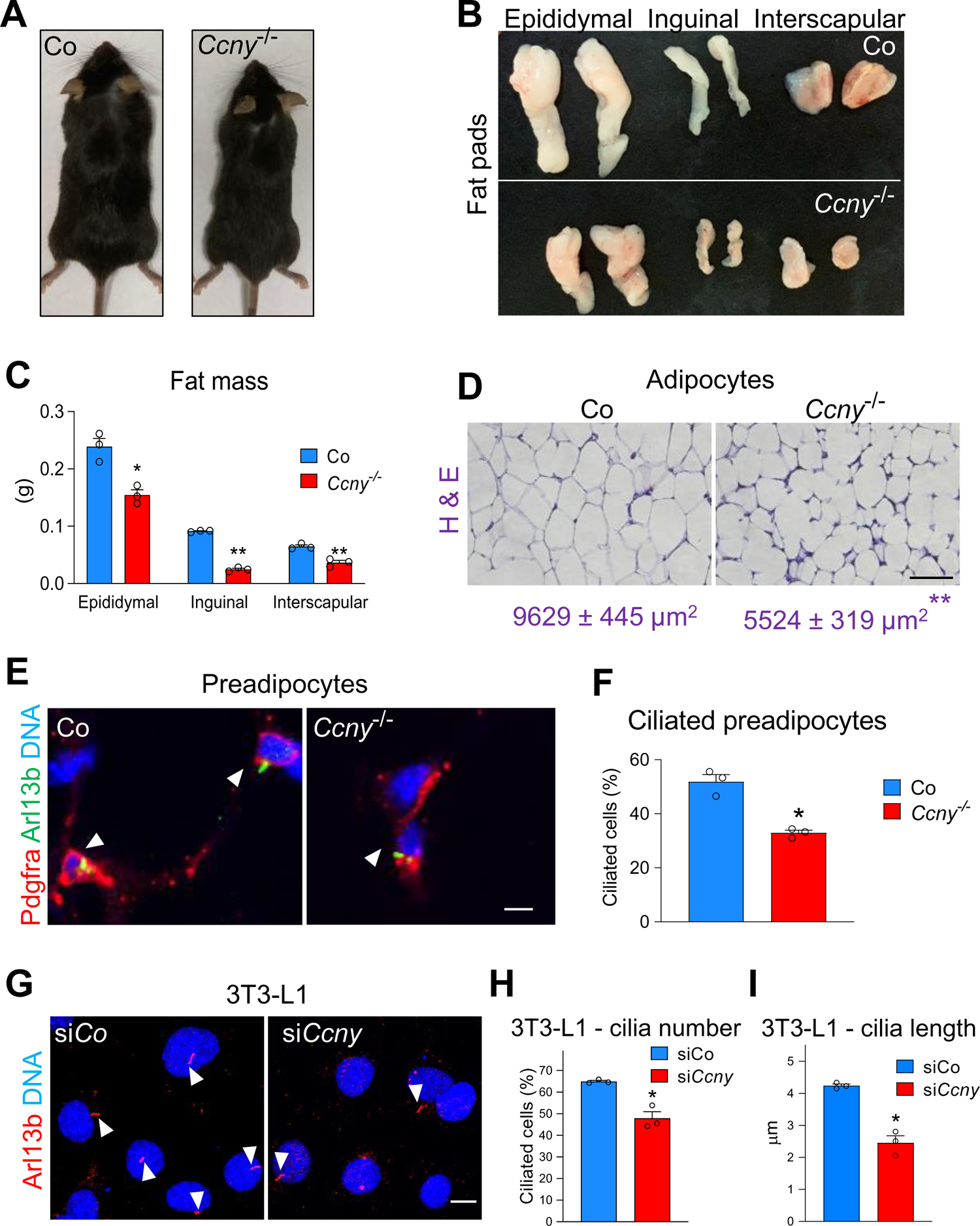
Lipodystrophy in *Ccny^-/-^* Mice due to Reduced Ciliogenesis. **(A)** Representative image of male *Ccny^-/-^* mutant mouse compared to control (*Ccny*^+/-^) littermate. Mutant mouse exhibits smaller body size. Scale bar 10 mm. **(B)** Comparison of epididymal, inguinal and interscapular fat pads between control and *Ccny^-/-^* adult male mice. All fat pads shown are from the same picture. **(C)** Quantification of epididymal, inguinal and interscapular fat pad weight (measured in grams (g)) from control and *Ccny^-/-^* mice. **(D)** H&E staining on paraffin sections from control and *Ccny^-/-^* epididymal fat pads. Average size of adipocytes is significantly decreased in *Ccny^-/-^* mice. Numbers (purple) are means of adipocyte size ± SEM (n=3 mice analyzed). Asterisks denote statistical significance. Scale bar 15 µm. **(E)** Co-IF for Arl13b and the preadipocyte marker Pdgfrα (platelet-derived growth factor receptor alpha) on sections of epididymal fat pads. White arrowheads depict ciliated preadipocytes. Scale bar 5 µm. **(F)** Quantification of ciliated preadipocytes from **(E)**. Data are means ± SEM (n=3 mice analyzed). **(G)** Acute siRNA knockdown of *Ccny* in mouse 3T3-L1 preadipocytes followed by IF for Arl13b to identify primary cilia. Cilia marked by white arrowheads. Scale bar 5 µm. (H-I) Quantification of cilia number **(H)** and length **(I)** in 3T3-L1 preadipocytes after siRNA knockdown of *Ccny*. Data are means ± SEM (experiment performed 3 times in triplicates, representative experiment shown). Data information: Unpaired two-tailed t-test for all statistical analyses: ns, not significant; * p < 0.05, ** p < 0.01.

### CCNY/L1 Promote Ciliogenesis through β-catenin-independent WNT/GSK3 Signalling

To dissect the molecular mechanisms underlying the role of *CCNY/L1* in primary cilia formation, we turned to HEK293T (293T) cells, a model system to study ciliogenesis mechanistically (Paridaen et al., 2013). Using CRISPR/Cas9 genome editing, we generated 293T cells deficient for *CCNY* and *CCNYL1* (referred to as *CCNY/L1*DKO cells; Figure S2A, schematic). Immunoblot analysis confirmed that CCNY and CCNYL1 proteins were not translated in *CCNY/L1* individual KO and DKO cells (Figure S2B). CCNY/L1 are M-phase cyclins and their hallmark in dividing cells is that they prime and sensitize LRP6 predominantly during G2/M phase (Davidson et al., 2009). Consequently, when HEK293T cells were arrested in G2/M with nocodazole they showed a 52% increase in SuperTOPFLASH reporter assays in response to WNT3A stimulation compared to non-synchronized cells. By contrast, in DKO cells, the WNT3A-stimulated increase in canonical WNT signalling during G2/M was abolished (Figure S2C). These results are consistent with CCNY/L1 predominantly mediating mitotic WNT signalling (Davidson et al., 2009).

Ciliogenesis can be induced in 293T cells by cell cycle exit upon serum depletion (Paridaen et al., 2013). Hence, we serum starved 293T cells and performed IF for ARL13B on the cell cycle-arrested cells, which revealed in *CCNY/L1*DKO cells a drastic decrease in cilia number (−74%, p = 0.0002) and length (−67%, p = 0.0007) (Figure 3A-C). To confirm that *CCNY/L1* regulation of primary cilia levels in 293T cells was related to WNT signalling, we analyzed a previously characterized *LRP5/LRP6*-deficient (*LRP5/6*DKO) 293T cell line (Kirsch et al., 2017). In *LRP5/6*DKO cells there was a similar reduction in cilia number (−65%, p = 0.0006) and length (−65%, p = 0.003) (Figure 3A-C).

**Figure 3.**
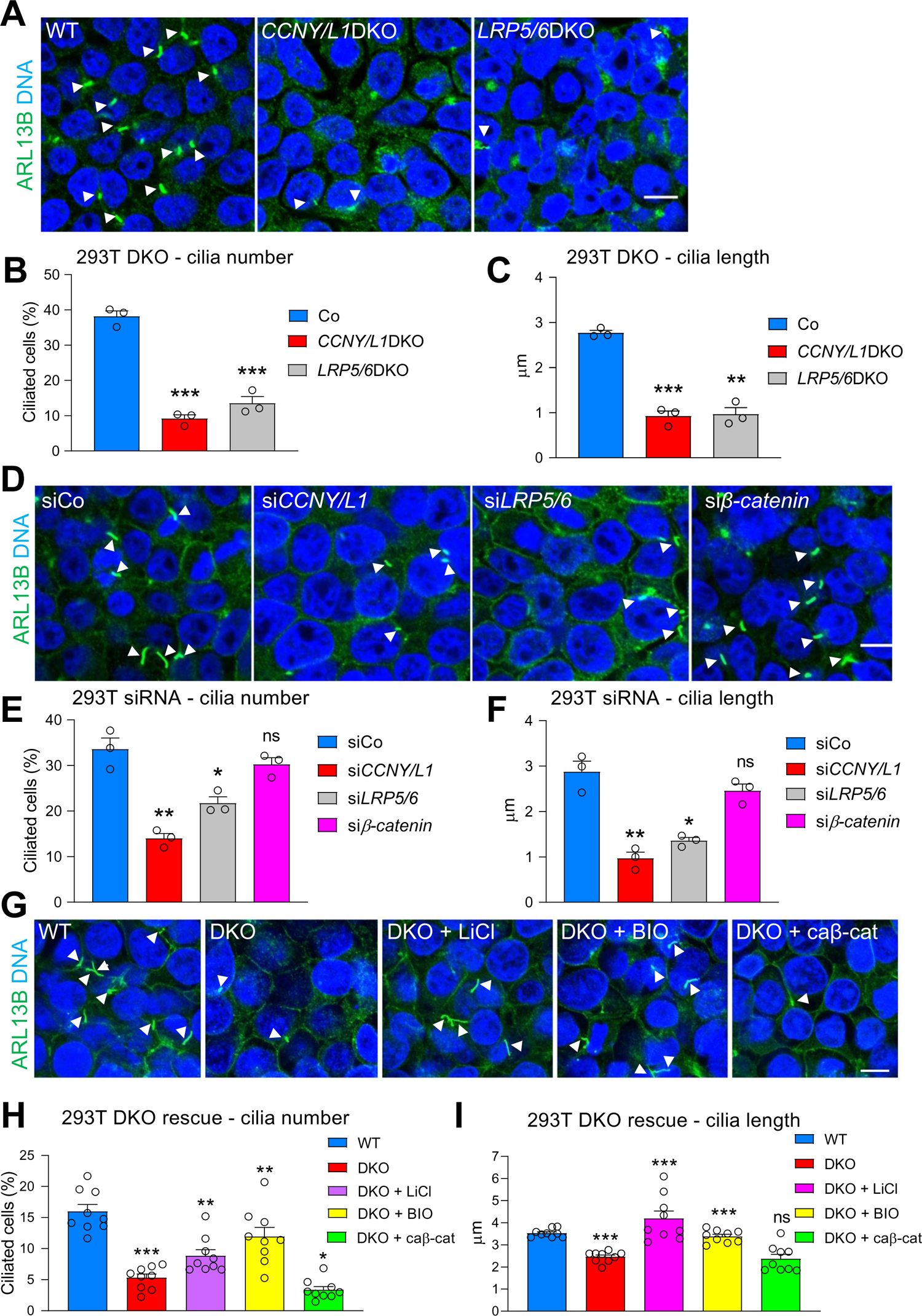
CCNY/L1 Promote Primary Cilia Formation through β-catenin-independent WNT/GSK3 Signalling. (**A**) ARL13B IF in serum-starved wildtype (WT), *CCNY*/*CCNYL1* (*CCNY/L1*) DKO and *LRP5/LRP6* (*LRP5/6*) DKO HEK293T (293T) cells to identify primary cilia (white arrowheads). (**B-C**) Quantification of the number **(B)** and length **(C)** of cilia in WT and DKO cell lines. Data are means ± SEM (experiment performed 3 times in triplicates, representative experiment shown). (**D**) Acute siRNA knockdown of *CCNY/L1*, *LRP5/6* and *β-catenin* followed by ARL13B IF to identify primary cilia (white arrowheads) in 293T cells. (**E-F**) Quantification of cilia number (E) and length (F) from (D). Data are means ± SEM (experiment performed three times in triplicates, representative experiment shown). (**G**) Twenty four-hour treatment with the GSK3 inhibitors Lithium Chloride (LiCl) (10 mM) or BIO (1 µM) and transfection of constitutively active *Xenopus* ΔN-β-catenin plasmid (caβ-cat) in wildtype and *CCNY/L1* DKO cells followed by ARL13B IF to identify primary cilia (white arrowheads). (**H-I**) Quantification of cilia number (H) and length (I) from (G). For statistical analyses, *CCNY/L1* DKO cells compared to wildtype, and pharmacological treatments/caβ-cat transfection compared to DKO cells. Data are means ± SEM (experiment performed 3 times in triplicates, all samples included in analysis). Data information: All scale bars 5 μm. Unpaired two-tailed t-test for all statistical analyses: ns, not significant; * p < 0.05, ** p < 0.01, *** p < 0.001.

Reduced ciliogenesis was not due to general cell cycle defects (e.g., mitotic arrest) since both *CCNY/L1* and *LRP5/6*DKO cells exited the cell cycle (Ki67⁻) in response to serum starvation at comparable rates to controls (Figure S2D-E). To support that cilia defects were not due to off-target effects from CRISPR/Cas9, we performed acute knockdown of *CCNY/L1* and *LRP5/6* using siRNA, which also led to reduced cilia number and length (Figure 3D-F). In contrast, siRNA knockdown of *β-catenin* did not lead to cilia defects (Figure 3D-F), despite high knockdown efficiency (Figure S2F). We performed rescue experiments in *CCNY/L1*DKO cells using different modes of WNT pathway stimulation. Treatment with the GSK3 inhibitors lithium chloride or BIO (6-bromoindirubin-3-oxime) increased the number and length of cilia in DKO cells, while overexpression of constitutively active β-catenin had no effect (Figure 3G-I). All treatments activated canonical WNT signalling to similar levels, as shown by SuperTOPFLASH assay and nuclear β-catenin accumulation (Figure S2G-H). In summary, these data corroborate that in non-dividing cells, *CCNY/L1* promote ciliogenesis via β-catenin-independent WNT/GSK3 signalling.

Previous work showed that LRP5/6 deficiency does not lead to cilia defects in Retinal Pigment Epithelium 1 (RPE1) cells, as well as HEK293 cells, the parent line of HEK293T cells (Bernatik et al., 2021). To confirm this finding, we performed siRNA knockdown of *LRP5/6* in RPE1 and HEK293 cells and indeed observed no significant differences in cilia number in both cell lines (Figure S2I-K). These results suggest that WNT/GSK3 signalling does not promote ciliogenesis universally and, instead, indicate context dependence.

### WNT Signalling Components Localize to Primary Cilia

Primary cilia are physically separated from the rest of the cell cortex by a selective barrier, the transition zone, which regulates the entry and exit of specific proteins (Garcia-Gonzalo and Reiter 2017; Gonçalves and Pelletier 2017; Nachury 2018). This separation also applies to the plasma membrane in that the constituents of the ciliary membrane are distinct from those in the planar plasma membrane and other plasma membrane protrusions such as microvilli. As a result, primary cilia contain a distinct protein composition and their membranes concentrate receptors of selected signalling pathways (Schneider et al., 2005; Rohatgi et al., 2007). Given the conserved role of WNT/GSK3 signalling in promoting ciliogenesis, we therefore addressed or confirmed the presence of WNT/GSK3 signalling components in primary cilia.

In serum-starved 293T cells, IF revealed highly specific ciliary localization of endogenous CCNY in 95% of cilia analyzed (n=35) (Figure 4A). CCNYL1 also localized to cilia (Figure 4B), although at a lower frequency (50%, n=30). CCNY/L1 staining was absent in *CCNY/L1*DKO cells, confirming specificity of the antibodies (Figure S3A-B). By IF, we confirmed GSK3β localization in primary cilia (Figure 4C), as previously reported (Thoma et al., 2007). The GSK3β antibody was validated by siRNA knockdown of *GSK3β* (Figure S3C). The Wnt receptor Frizzled 5 (FZD5) was also detected in primary cilia by IF upon overexpression (Figure 4D), demonstrating that FZD receptors can localize to cilia in 293T cells, as previously shown in other cell lines (Stypulkowski et al., 2021).

**Figure 4.**
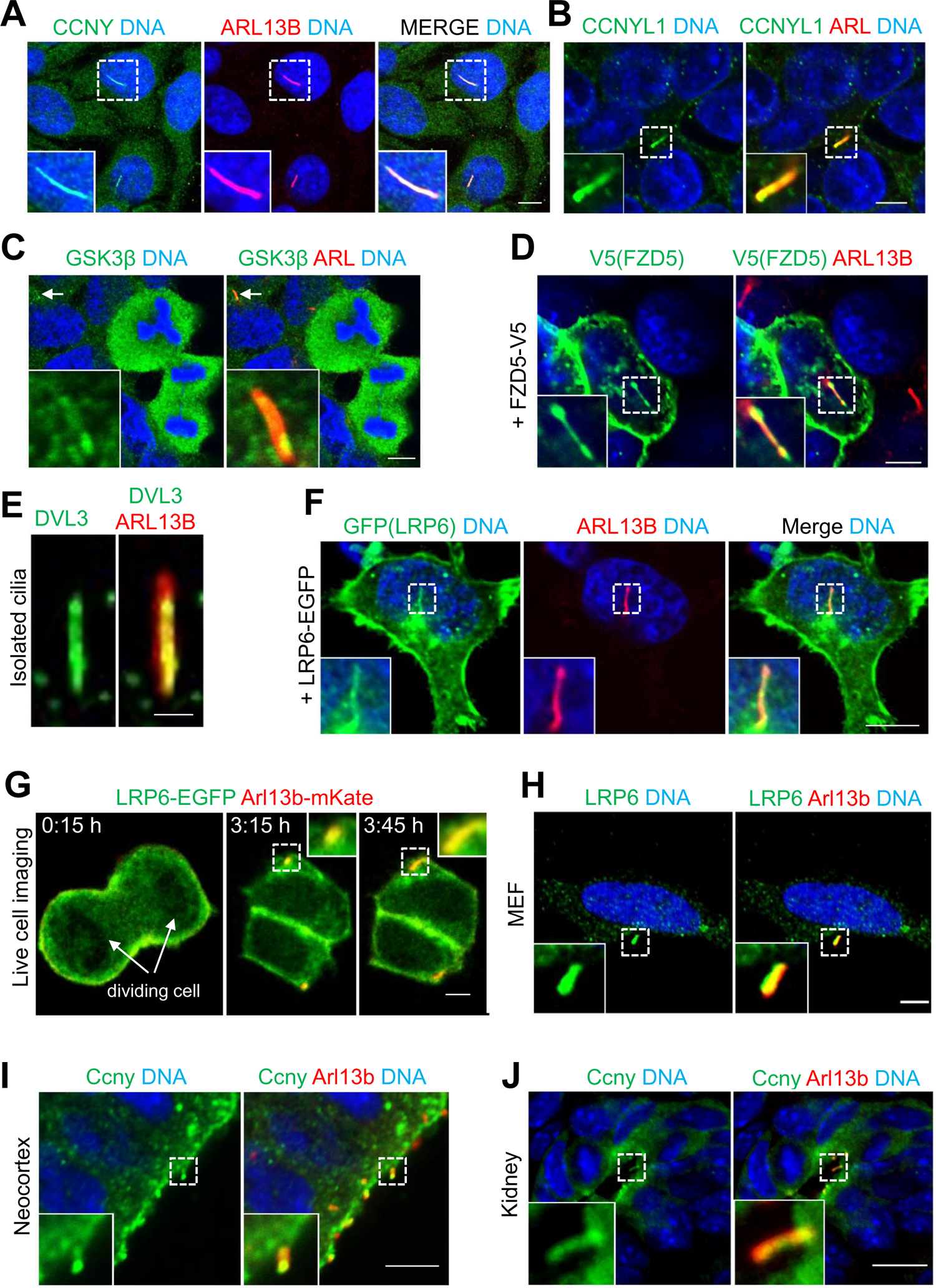
WNT Signalling Components Localize to Primary Cilia. **(A)** Co-IF for CCNY and ARL13B in 293T cells. Scale bar 5 µm. **(B)** Co-IF for CCNYL1 and ARL13B **(ARL)** in 293T cells. Scale bar 5 µm. **(C)** Co-IF for GSK3β and ARL13B in 293T cells. Cilium indicated by arrow and magnified in inset. Scale bar 5 µm. **(D)** Co-IF for V5 tag and ARL13B in 293T cells transfected with FZD5-V5 plasmid. Scale bar 5 µm. **(E)** Co-IF for DVL3 and ARL13B on cilia isolated from 293T cells using poly-D-lysine coated cover slips. Scale bar 1 µm. **(F)** Co-IF for GFP and ARL13B in 293T cells transfected with LRP6-EGFP plasmid. Scale bar 5 µm. **(G)** Still images acquired during live cell imaging of a 293T cell upon exiting mitosis (cell division shown at 0:15 h) and forming cilia. Cells were transfected with LRP6-EGFP (green) and pArl13b-mKate2 (red) plasmids. Cells were shifted to serum-free medium at t = 0. Note co-staining of LRP6 and ARL13B (yellow) in nascent cilium (inset at 3:30 h) and elongated cilium (inset at 4:00 h). Scale bar 2.5 µm. **(H)** Co-IF for endogenous LRP6 and Arl13b in mouse embryonic fibroblast (MEF) cells. Scale bar 5 µm. **(I)** Ccny and Arl13b co-IF in the neocortex of E13.5 embryos reveals Ccny localization to apical primary cilia (magnified in inset). Scale bar 10 µm. **(J)** Ccny is detected in primary cilia of developing kidney proximal tubules (white arrowhead, magnified in inset). Scale bar 10 µm. Data information: Where indicated white dashed boxes magnified in insets. IF analysis on cell lines performed after 24 hours of serum starvation.

Dishevelled (DVL) proteins are key components of the WNT pathway that relay WNT signals from receptors to downstream effectors (Niehrs, 2012; Jeong and Jho, 2021). Moreover, DVLs have been previously shown to localize to cilia (Veland et al., 2013; Gupta et al., 2015). To check for the presence of DVL proteins in 293T primary cilia, we performed IF analysis on serum-starved cells with antibodies against DVL1, 2, or 3. Of the 3 proteins, only DVL3 was detected in primary cilia in 293T cells (antibody validated by siRNA, see Figure S3D; data for DVL1 and 2 not shown). To ensure ciliary DVL3 staining was not due to superimposition of cytoplasmic signals, we isolated cilia using poly-D-lysine coated cover slips (Huang et al., 2009) and then performed IF for DVL3, which revealed highly specific ciliary staining (Figure 4E). To test if DVL3 is required for ciliogenesis in 293T cells, we performed siRNA knockdown of *DVL3*, which led to reduced cilia number (Figure S3E). Triple siRNA knockdown of *DVL1/2/3* led to a slight but significant further reduction in cilia number when compared to si*DVL3* alone (Figure S3F), suggesting DVL1 or 2 may still localize to cilia in 293T cells but the levels are too low to be detected with the antibodies used. Nevetheless, individual knockdown of *DVL1* or *DVL2* in both cases did not lead to cilia defects (Figure S3G), altogether suggesting that DVL3 is the principle family member involved in primary cilia formation in 293T cells, while DVL1 and 2 likely only play minor roles. Importantly, the si*DVL1/2/3-*induced cilia defect was rescued by treatment with the GSK3 inhibitor CHIR99021, confirming that DVLs regulate ciliogenesis in 293T cells through WNT/GSK3 signalling (Figure S3H).

We next analyzed the localization of the WNT co-receptor LRP6. Since *LRP6* expression levels in 293T cells are too low to be detected with available antibodies, we overexpressed LRP6-EGFP and found it localized to the ciliary membrane in 59% of cells analyzed (n=46) (Figure 4F). This effect was unrelated to the EGFP moiety since we also detected LRP6 in cilia upon transfection of non-tagged LRP6 in 65% of cells analyzed (n=34) (Figure S3I).

We carried out live cell imaging of 293T cells co-transfected with LRP6-EGFP and pArl13b-mKate2 to examine the kinetics of LRP6 localizing to cilia upon serum starvation. Close inspection showed that when the ARL13B punctate cilia rudiment first appeared as early as 3 hours after cell division, it immediately co-localized with LRP6, suggesting an intimate link from the very onset of ciliogenesis (Figure 4G; Video S1). Immediate co-localization of LRP6 with ARL13B in developing cilia was observed in 18 out of 20 newly-divided cells.

To confirm ciliary localization of endogenous LRP6, we analyzed serum-starved mouse embryonic fibroblasts (MEFs), which express higher levels of *Lrp6*. Consistently, 30% (n=100) of MEF cilia displayed highly specific LRP6 localization, demonstrating that endogenous LRP6 localizes to the ciliary membrane (Figure 4H). LRP6 staining was absent in *LRP5/6*DKO MEFs, validating the IF staining (Figure S3J). Confirming ciliary localization *in vivo*, IF for Ccny and Arl13b in the neocortex and kidney proximal tubules showed prominent co-localization in both tissues (Figure 4I-J). In summary, we confirm and extend previous studies that key components of the WNT/GSK3 cascade localize to primary cilia in both human and mouse cells.

### Identification of a Ciliary Targeting Sequence in LRP6

Large ciliary transmembrane proteins (e.g. > 40 kDa) frequently contain ciliary targeting sequences (CTS) that direct them to cilia and facilitate their entry through the transition zone (Malicki and Avidor-Reiss, 2014). Two of the most common CTSs identified are the VxP (VP) motif, typically located in the C-terminus (Mazelova et al., 2008; Higginbotham et al., 2012), and the AxxxQ (AQ) motif, which features in several GPCRs (Berbari et al., 2008). Inspection of human LRP6, a large transmembrane protein (>200 kDa), revealed a C-terminal VP motif (aa 1567-1569, <u>V</u>P<u>P</u>) (Figure S4A) and an N-terminal AQ motif (aa 272-276, <u>A</u>FSQ<u>Q</u>) (Figure 5A). Both motifs are highly conserved in mammals. We performed site-directed mutagenesis of both putative LRP6 CTSs (see Figure S4A and Figure 5A), transfected the mutated constructs into 293T cells, and induced ciliogenesis by serum starvation. Strikingly, mutating the AQ motif greatly reduced (−85%, p=0.0004) ciliary LRP6 localization (LRP6**^+^** cilia) (Figure 5B-C), while the VP mutant showed only a slight but non-significant reduction (−23%, p=0.06) (Figure S4B). Double mutation of the VP and AQ motifs did not lead to a further reduction of LRP6**^+^** cilia compared to the LRP6^AQ>F^ mutant alone, suggesting AQ to be the principle CTS for LRP6 (Figure S4B). Interestingly, the AQ motif was not conserved in LRP5 and transfected Myc-tagged LRP5 did not show any ciliary localization (0 out of 33 cilia analyzed) (Figure 5D-E), further corroborating the importance of the AQ motif for LRP6 ciliary targeting. We conclude that the principle WNT/GSK3 receptor, LRP6, contains an ‘AQ’ CTS in its N-terminus, supporting that its ciliary localization is an evolved function.

**Figure 5.**
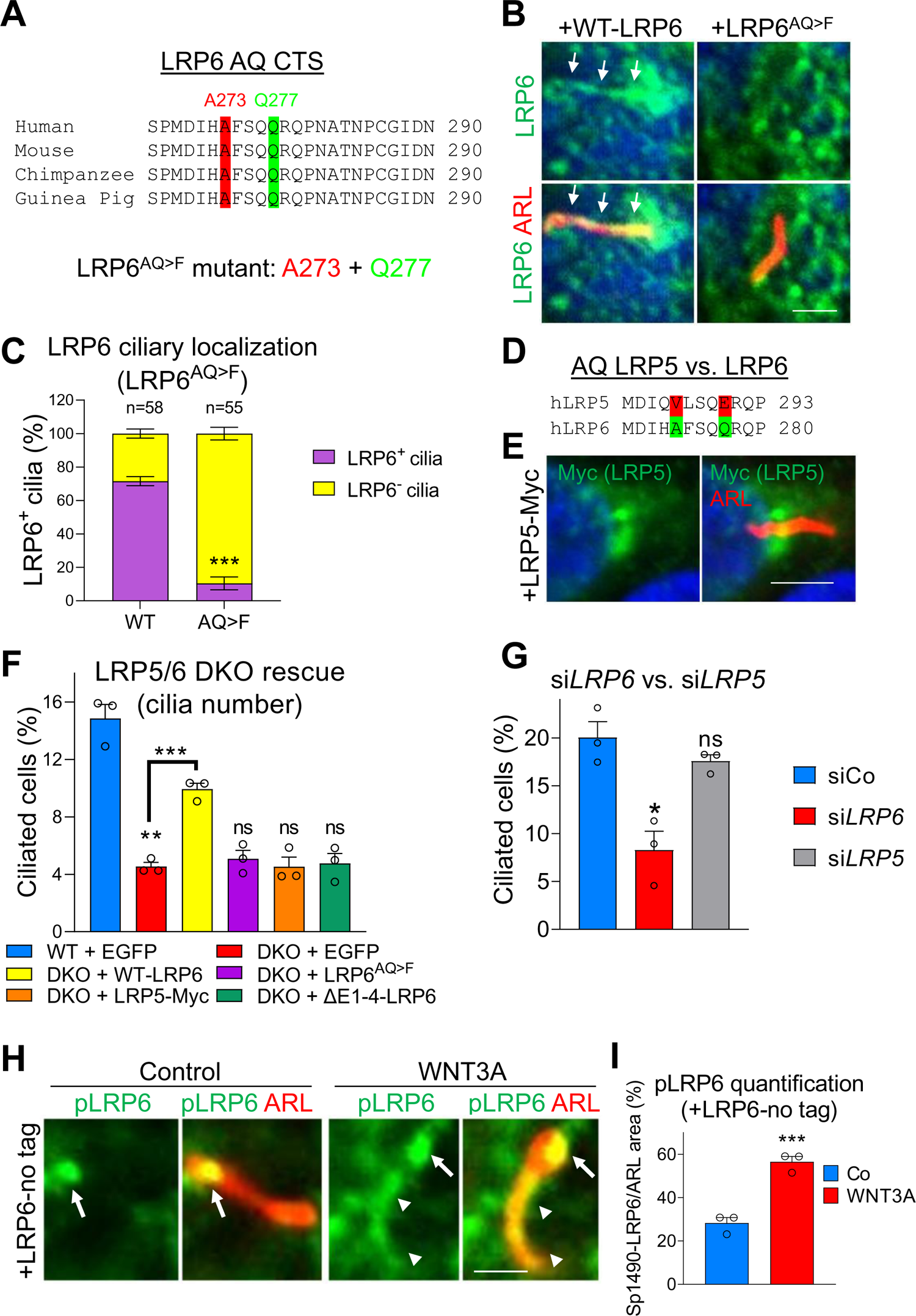
Primary Cilia are WNT Transducing Organelles. **(A)** Amino acid sequence alignment of LRP6 from human, mouse, chimapanzee and guinea pig showing the conserved AQ ciliary targeting sequence (CTS). The AQ-mutant of LRP6 was generated by mutating both alanine **(A)** and glutamine (Q) to phenylalanine **(F)**. **(B)** Transfection of WT-LRP6 or LRP6^AQ>F^ mutant into 293T cells followed by 24 hour serum starvation and co-IF for LRP6 and ARL13B. White arrows depict LRP6 ciliary staining. Scale bar 2 µm. **(C)** Quantification of **(B)**. Data are means ± SEM. n = number of transfected cells with cilia, identified by ARL13B IF. Experiment performed 3x in duplicates and results pooled for final analysis. **(D)** Amino acid sequence alignment of human LRP5 with human LRP6. The AQ LRP6 CTS is not conserved in LRP5. **(E)** Co-IF for Myc and ARL13B in 293T cells transfected with Myc-tagged LRP5 plasmid and serum starved for 24 hours. LRP5 is not detected in primary cilia. DNA shown in blue. Scale bar 2.5 μm. **(F)** Cilia rescue experiment in *LRP5/6*DKO 293T cells. Cells were transfected with indicated plasmids for 24 hours, and then serum starved for an additional 24 hours. EGFP plasmid was used as control for WT and DKO. Cilia number was monitored by IF for ARL13B. Data are means ± SEM. For statistical analyses, DKO compared to WT, and DKO + plasmids compared to DKO. Experiment performed twice in triplicates, representative experiment shown. **(G)** Acute siRNA knockdown of LRP6 or LRP5 in 293T cells followed by serum starvation and quantification of the number of cilia (identifed by IF for ARL13B). Data are means ± SEM (experiment performed twice in triplicates, representative experiment shown). **(H)** IF for phospho-LRP6 (Sp1490) and ARL13B in 293T cells transfected with non-tagged LRP6, serum-starved for 48 hours, and treated with WNT3A conditioned media for 30 minutes. Arrows depict phospho-LRP6 staining in ciliary tips while arrowheads depict staining throughout the entire ciliary shaft. Scale bar 2 µm. **(I)** Quantification of **(H)**. Phospho-LRP6 (Sp1490) pixel area was normalized to ARL13B pixel area. Data are means ± SEM (experiment performed twice in triplicates, representative samples shown). Data information: Unpaired two-tailed t-test for all statistical analyses: ns, not significant; * p < 0.05, ** p < 0.01, *** p < 0.001.

### Primary Cilia are WNT Transducing Organelles

Identification of the LRP6 CTS provided the opportunity to address directly the question whether primary cilia are WNT-signalling organelles. Previous reports as well as the data presented thus far fail to answer whether WNT-driven ciliogenesis is due to WNT signalling happening within cilia or rather proceeding through the non-ciliary regions of the plasma membrane followed by cytoplasmic relay to cilia. To uncouple cytoplasmic from ciliary WNT signalling, we conducted rescue assays in cilia-deficient *LRP5/6*DKO 293T cells. If ciliary WNT signalling proceeds at- and within the cilium, then only WT-LRP6 but neither the LRP6^AQ>F^ mutant, nor LRP5, nor constitutively active ΔE1-4-LRP6 (Mao et al., 2001), that all lack a CTS, should rescue ciliogenesis. As a control, SuperTOPFLASH assay in *LRP5/6*DKO cells confirmed that the LRP6^AQ>F^ and ΔE1-4-LRP6 mutants transduced canonical WNT signalling at comparable or even higher levels compared to WT-LRP6, while LRP5-Myc, in accordance with previously published data (MacDonald et al., 2011), displayed 50% less WNT activation (Figure S4C). To compensate for the latter, we transfected twice the amount of LRP5-Myc plasmid. Transfection efficiencies were monitored by IF (Figure S4D). Expectedly, WT-LRP6 robustly rescued ciliogenesis in serum-starved *LRP5/6*DKO 293T cells (p=0.0009) (Figure 5F). Importantly, neither LRP6^AQ>F^ mutant, nor ΔE1-4-LRP6, nor LRP5-Myc were able to rescue the cilia defect, strongly corroborating that proper ciliogenesis requires ciliary-localized LRP6 and hence signalling within cilia. Finally, we performed single siRNA knockdown of either *LRP5* or *LRP6* in 293T cells. Since only LRP6 localizes to cilia, we reasoned that individual knockdown of *LRP6*, but not *LRP5*, should induce ciliogenesis defects. Consistently, si*LRP6* led to decreased cilia number while si*LRP5* had no effect (Figure 5G).

We next tested if cilia can directly respond to exogenous WNT stimulation by monitoring acute receptor phosphorylation. Primed and active, ligand-triggered WNT signalling can be monitored with an LRP6 antibody specific for the cyclin Y-CDK14/GSK3β phosphorylation site Sp1490 (Davidson et al., 2009; Zeng et al., 2005). This LRP6 phosphorylation-site is crucial because it sequesters and inhibits GSK3 (Piao et al., 2008). Following overexpression of non-tagged LRP6 in 293T cells, Sp1490-LRP6 was detected after 48 hrs serum starvation in cilia, where the signal tended to localize at the tips (Figure 5H), and hence may be related to cilia-derived ectosomes (Dubreuil et al., 2007; Wood and Rosenbaum, 2015). Strikingly, a 30 min WNT3A pulse led to a further increase in Sp1490-LRP6 ciliary staining (Figure 5H-I), demonstrating that 293T primary cilia respond to exogenous WNT ligands.

Taken together, we conclude that primary cilia respond directly to endogenous and exogenous WNTs, and that this response decomissions GSK3 to promote ciliogenesis, altogether demonstrating that primary cilia behave as WNT-signalling organelles.

### A WNT ┫ PP1 Signalling Axis Promotes Primary Cilia Formation

Since ciliary WNT/GSK3 signalling promotes cilia formation independent of β-catenin, what could be its downstream link to cilia formation instead? One candidate is serine/threonine protein phosphatase 1 (PP1), a well-established regulator of flagellar motility in *Paramecium*, *Chlamydomonas*, and mammalian sperm (Klumpp et al., 1990; Habermacher and Sale, 1996, Vijayaraghavan et al., 1996; Yang et al., 2000). Mechanistically, PP1 regulates phosphorylation and function of the intraflagellar transport motor KIF3B in *Chlamydomonas*, (Liang et al., 2018). PP1 also binds the ciliary ion channel polycystin-1 (PC1) and shortens the length of primary cilia in mammalian cells (Luo et al., 2019). PP1 is regulated by a large family of regulatory subunits, some of which are inhibitory and maintain the enzyme in an inactive state (Casamayor and Ariňo, 2020). Among the inhibitory subunits, PPP1R2 is inactivated by GSK3-dependent phosphorylation at Thr72 (Tp72), thereby activating PP1 (Cohen, 1989). Moreover, knockdown of PPP1R2 leads to reduced cilia number in RPE1 cells (Wang and Brautigan, 2008). We previously showed that during mouse sperm maturation, WNT signalling inhibits PP1 via the GSK3β ┫ PPP1R ┫ PP1 axis, leading to global increase in protein phosphorylation and unlocking of flagellar motility (Koch et al., 2015). Given the similarity between flagella and cilia, as well as the well-described roles of PP1 in regulating cilia formation, we hypothesized that WNT/GSK3 signalling may regulate ciliogenesis via PPP1R2-PP1 (Figure 6A, schematic).

**Figure 6.**
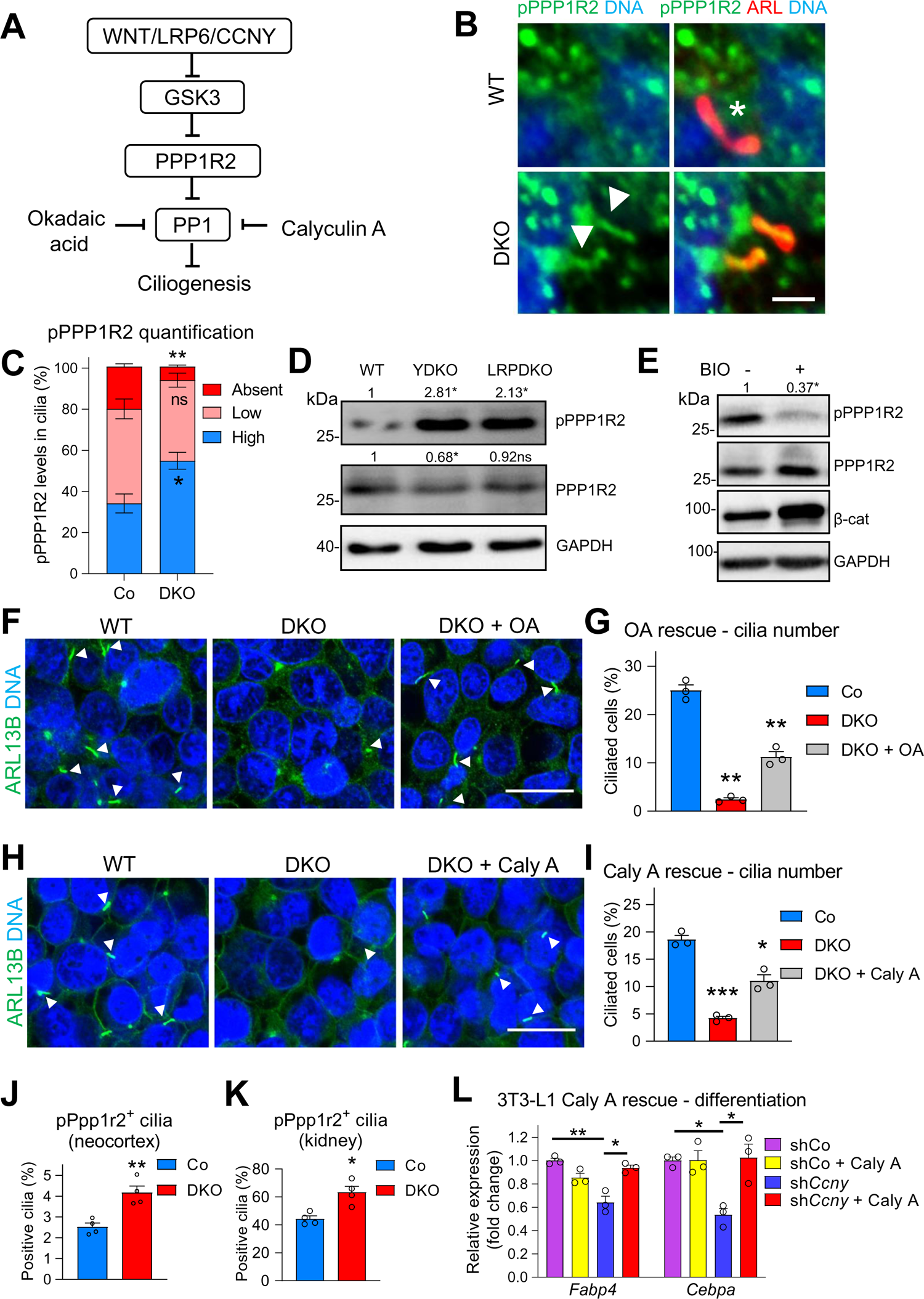
A WNT ┫ PP1 Signalling Axis Promotes Primary Cilia Formation. **(A)** Schematic of the post-transcriptional WNT/GSK3 signalling cascade that promotes primary cilia formation by inhibiting Protein phosphatase 1 (PP1). WNT/LRP6/CCNY signalling reduces inhibitory GSK3 phosphorylation of PPP1R2 and thereby inhibits PP1, a negative regulator of ciliogenesis. Okadaic acid **(OA)** and Calyculin A (Caly A) inhibit PP1 activity and rescue the effects of upstream WNT inhibition. **(B)** Co-IF for PPP1R2, phosphorylated at T72 (pPPP1R2), and ARL13B (ARL) in WT and *CCNY/L1* DKO HEK293T cells. Cilia with high (white arrowhead) or absent (white asterisks) pPPP1R2 staining indicated for WT and DKO cells. Scale bar 2.5 μm. **(C)** Quantification of pPPP1R2 ciliary staining from **(B)**. Data are means ± SEM (experiment performed twice in duplicates, all samples included in analysis). **(D)** Immunoblot analysis with pPPP1R2 antibody on *CCNY/L1*DKO (YDKO) and *LRP5/6*DKO (LRPDKO) whole cell lysates. Numbers above blots represent pPPP1R2 levels normalized to total PPP1R2 levels (top), and total PPP1R2 levels normalized to GAPDH (middle). GAPDH was used as a loading control. Asterisks depict statistical significance. **(E)** Immunoblot analysis on cell lysates from WT 293T cells treated with BIO (1 µM) overnight. Numbers above blot represent pPPP1R2 levels normalized to total PPP1R2 levels. Blot for total β-catenin (β-cat) confirms WNT pathway activation upon BIO treatment. GAPDH, loading control. **(F)** ARL13B IF on WT and DKO cells treated with 0.1 µM OA for 4 hours prior to serum starvation. Cilia depicted by white arrowheads. Scale bar 10 μm. **(G)** Quantification of cilia number from **(F)**. Cilia number in DKO cells compared to WT and OA treatment to DKO cells. Data are means ± SEM (experiment performed three times in triplicates, representative experiment shown). **(H)** ARL13B IF on WT and DKO cells treated with 0.5 nM Calyculin A (Caly A) during 24 hour serum starvation. Cilia depicted by white arrowheads. Scale bar 10 μm. **(I)** Quantification of cilia number from **(H)**. Cilia number in DKO cells compared to WT and Caly A treatment to DKO cells. Data are means ± SEM (experiment performed three times in triplicates, representative experiment shown). **(J)** Quantification of phospho-Ppp1r2^+^ (pPpp1r2^+^) cilia in the neocortex of E13.5 control and DKO embryos. Data are means ± SEM (n=3 embryos, 2 litters). Representative IF images shown in Figure S4L. **(K)** Quantification of phospho-Ppp1r2^+^ (pPpp1r2^+^) cilia in E13.5 control and DKO kidneys. Data are means ± SEM (n=3 embryos, 2 litters). Representative IF images shown in Figure S4M. **(L)** qPCR analysis on RNA extracted from 3T3-L1 preadipocytes transduced with sh*Ccny*-expressing lentivirus and induced to differentiate into mature adipocytes. Supplementation of differentiation media with Calyculin A (0.5 nM) rescues the expression of the adipocyte markers *Fabp4* and *Cebpa* in sh*Ccny*-treated cells. Data information: Unpaired two-tailed t-test for all statistical analyses: ns, not significant; * p < 0.05, ** p < 0.01, *** p < 0.001.

To test this hypothesis, we serum starved *CCNY/L1* and *LRP5/6*DKO 293T cells and performed IF with an antibody against the GSK3-specific Tp72 phosphorylation site of PPP1R2. The number of of Tp72-PPP1R2^+^ cilia as well as the staining intensity in individual cilia were both significantly increased in DKO cells (Figure 6B-C). This increase was not due to more PPP1R2 protein in cilia since IF for total PPP1R2 revealed similar staining intensity in DKO and control cilia (Figure S5A-B). Immunoblot analysis of cell lysates confirmed increased Tp72-PPP1R2 levels in *CCNY/L1* and *LRP5/6*DKO cells, indicating WNT-receptor engagement in regulation of PPP1R2 phosphorylation (Figure 6D). Conversely, BIO treatment drastically reduced PPP1R2 phosphorylation by immunoblot, corroborating PPP1R2 as a GSK3 substrate in 293T cells (Figure 6E).

Our models predict that ciliogenic WNT signalling functions to repress PP1 activity and hence that the ciliogenesis defects in *CCNY/L1*DKO cells are due to elevated PP1 activity. Hence, we carried out rescue experiments in *CCNY/L1*DKO cells using the general phosphatase inhibitors Okadaic acid (OA) and Calyculin A (Caly A) (Figure 6A, schematic). Consistenly, OA and Caly A treatment led to respective 4.6 (p=0.009) and 2.6 (p=0.02) fold inductions in cilia number in DKO cells (Figure 6F-I). Calyculin A treatment also moderately rescued cilia number in si*DVL1/2/3*-treated cells (Figure S5C). To confirm a functional role for PPP1R2 in primary cilia formation in 293T cells, we performed siRNA knockdown of PPP1R2 (knockdown efficiency confirmed in Figure S5D), which reduced cilia number and length (Figure S5E-G). Taken together, these results support a WNT ┫ PP1 axis in promoting ciliogenesis in 293T cells.

We sought to corroborate the presence of a ciliary WNT ┫ PP1 axis in a physiological context. Ppp1r2 was detected in primary cilia of the neocortex and kidney proximal tubules by IF (Figure S5H-I). In *Ccny/l1* DKO mouse embryos, co-IF for Tp72-Ppp1r2 and Arl13b in the neocortex and kidney showed an increased number of Tp72-Ppp1r2**^+^** primary cilia in the neocortex (+65%, p=0.007) and kidney proximal tubules (+43%, p=0.005) (Figure 6J-K; S5J-K). In 3T3-L1 cells we tested if the observed cilia and differentiation defects could be rescued by PP1 inhibition and found that in 3T3-L1 cells transduced with a sh*Ccny*-expressing lentivirus, Caly A treatment increased both cilia number (p=0.001) (Figure S5L) and the expression of the mature adipocyte markers *Fabp4* and *Cebpa* (Figure 6L). The results support that a WNT ┫ PP1 axis promotes ciliogenesis also *in vivo*.

## DISCUSSION

The three key findings of this study are i) that primary cilia are WNT signalling organelles that transduce a WNT response distinct from that of the plasma membrane, ii) that this response triggers a WNT ┫ PP1 axis, which promotes formation of primary cilia, and iii) that deficiency in this WNT signalling axis recapitulates ciliopathies. These conclusions are supported in a variety of biological model systems, including human cell lines, mouse embryos, and adult mice. We monitor acute (phospho-) LRP6 signalling and identify and mutate a ciliary localization signal in LRP6 to uncouple plasma membrane from ciliary signal transmission. We conclude that an evolutionary conserved WNT ┫ PP1 signalling axis acts as a common regulator of primary cilia formation in multiple cell types.

### Primary Cilia as WNT Transducing Organelles

One main conclusion of this study is that primary cilia are genuine WNT transducing organelles that signal via the WNT/GSK3 signalling machinery. While this WNT response employs upstream components of the canonical WNT pathway, it is independent of β-catenin. Instead, WNT activation of the LRP6 co-receptor decommissions GSK3 to inactivate an inhibitor of ciliogenesis, PP1.

Which steps of the WNT signalling cascade proceed in cilia? Generally, WNT ligands trigger the high affinity WNT receptor FZD and the co-receptor LRP6 to form a ternary complex. FZD5 and −6 were identified in the human cilia proteome (Gupta et al., 2015) and FZD6 localizes to the ciliary axoneme of HEK293 cells (Strakova et al., 2018). Consistently, we have detected FZD5 in the ciliary axoneme by overexpression in 293T cells. Signalling is routed to the canonical pathway via the WNT co-receptor LRP6 that we show carries an ‘AQ’ CTS to translocate the protein to the ciliary membrane. Specifically, live cell imaging of LRP6 in serum-starved cells shows that LRP6 co-localizes with ARL13B as soon as the latter accumulates at the nascent cilium base.

LRP6 signalling requires DVL polymers, platforms that cluster WNT receptor complexes into LRP6 signalosomes (Bilic et al., 2007). All three DVL proteins are prominent ciliary components, either in the centriole or the axoneme (Gupta et al., 2015) and they are required for ciliogenesis (Park et al., 2008; Lee et al., 2012; Ohata et al., 2014). Consistently, we show that DVL3 localizes to cilia in 293T cells, and that siRNA knockdown of DVL3 induces ciliogenesis defects, similar to acute *CCNY/L1* and *LRP5/6* deficiency. LRP6 becomes phosphorylated by CK1γ at T1479 (Davidson et al., 2005) and this phosphorylation is phospho-primed via mitotic CCNY/CDK14/16 at nearby PPPS/TP motifs (e.g. pS1490) (Davidson et al., 2009). The observation of ciliary LRP6 phosphorylation and its stimulation by WNT treatment is significant for two reasons: First, it indicates that all essential events leading up to signalosome formation proceed in the cilium. Second, it signifies GSK3 inhibition since the S1490 phospho-site sequesters and inhibits GSK3 (Piao et al., 2008). This GSK3 inhibition prevents PPP1R2 from becoming phosphorylated and inactivated, allowing it to repress PP1. Since ciliary PPP1R2 phosphorylation is reduced in *CCNY/L1* and *LRP5/6* deficient 293T cells we conclude that the entire WNT ┫ PP1 signalling axis is active in primary cilia.

Ciliary WNT signalling as reported here does not exclude the possibility of signal transmission through cilia into the cytoplasm. Since β-catenin can localize to the axoneme (Corbit et al., 2008) it might be modulated by ciliary WNT signalling and shuttle back into the cytoplasm. What the effects of ciliary WNT signalling are on β-catenin and how ciliary WNT signalling relates to the contentious question whether cilia restrain WNT/β-catenin signalling, which we have not addressed in this study, could be explored in the future. In this context, it is interesting that hedgehog signalling triggers distinct ciliary- and cytoplasmic reponses (Truong et al., 2021). Moreover, ciliary WNT signalling is part of a multipronged WNT-driven ciliogensis program since transcriptional WNT β-catenin signalling promotes cilia formation also transcriptionally (Caron et al., 2012; Walentek et al., 2012; Li Song et al., 2014; Walentek et al., 2015; Walentek et al., 2015b; Kyun et al., 2020; Zhang et al., 2020).

Mitosis is the stage when cilia disassemble during the cell cycle (Rieder et al., 1979; Tucker et al., 1979; Ehler et al., 1995; Wheatley et al., 1996; Paridaen et al., 2013). Hence, it seems paradoxical that mitotic cyclins CCNY/L1 concentrate in the axoneme and are required for ciliogenesis of quiescent cells (e.g. serum-starved cell lines, mouse spermatocytes). However, our finding is in keeping with accumulating evidence that multiple cyclins and CDKs are redeployed in quiescent cells to regulate motile ciliogenesis without affecting cell cycle progression (Tam et al., 2007; Mikolcevic et al., 2012; Wallmeier et al., 2014; Al Jord et el., 2017; Vladar et al., 2018). This redeployment of *bona fide* cell cycle regulators may reflect commonalities in the regulation of centrioles during ciliogenesis and the cell cycle (Al Jord et al., 2019).

### A WNT ┫ PP1 Axis Promotes Ciliogenesis

The second important finding of this study is the elucidation of a WNT ┫ PP1 signalling axis as the underlying mode whereby WNT promotes ciliogenesis in primary cilia. Yet, not all ciliated cells require WNT signalling for ciliogenesis, such as RPE1 and HEK293 cells (Bernatik et al., 2021 and this study), suggesting that the effects of WNT signalling on ciliogenesis are not universal. Even in cells that do engage ciliary WNT/GSK3 signalling, the requirement is not absolute since in *LRP5/6DKO* 293T cells, and in *Ccny/l1* DKO mice, ciliogenesis was reduced but not abolished. The reasons for the WNT-independence in certain cell types are unclear but may be due to alternative modes of PP1 regulation, known to be subject to diverse inputs (Brautigan and Shenolikar, 2018). Altogether, these data indicate a modulatory and context-dependent role of ciliary WNT/GSK3 signalling.

At the heart of the ciliogenic WNT cascade lies GSK3 inhibition. Consistent with this conclusion, this ubiquitous kinase localizes to cilia and is a well-established negative regulator of ciliogenesis that is evolutionary conserved: In *Chlamydomonas*, zebrafish embryos, and mammalian cells, GSK3 inhibition is well known to induce cilium elongation (Wilson and Lefebvre, 2004; Miyoshi et al., 2009; Ou et al., 2012; Yuan et al., 2012; Han et al., 2014; Nakakura et al., 2015; Thompson et al., 2016). Yet, GSK3β can also promote ciliogenesis e.g. by regulating the ciliary effectors Dzip1 and pVHL (Thoma et al., 2007; Zhang et al., 2015). The difference may be due to context- or dose dependence since e.g. in *Chlamydomonas* pharmacological inhibition of GSK3 for one hour induces long cilia while longer treatment induces an aflagellate phenotype (Wilson and Lefebvre, 2004). It is thought that GSK3 can reside in different pools (Thoma et al., 2007; Zhang et al., 2015), possibly biomolecular condensates (Schaefer and Peifer, 2019), which may also have distinct roles during ciliogenesis. Moreover, GSK3β activity can also be inhibited by Insulin and Insulin-like growth factor signalling, both of which can trigger IGFR/AKT signalling in cilia (Christensen et al., 2017). This suggests that the ciliary GSK3 ┫ PPP1R ┫ PP1 axis may also be regulated by other growth factors.

We identify the PP1 inhibitor PPP1R2 as a key target of the WNT/GSK3 module. PPP1R2 is concentrated in cilia and is a known effector of primary cilium- and flagellum morphogenesis upstream of PP1 (Cohen, 1989; Wang and Brautigan, 2008). Mechanistically, PP1 binds polycystin-1 (PC1) and shortens the length of primary cilia in mammalian cells (Luo et al., 2019). Furthermore, PP1 regulates the intraflagellar transport (IFT) system via phosphorylation of the kinesin motor protein FLA8/KIF3B in *Chlamydomonas* (Liang et al., 2018). Since WNT signalling induces a dramatic increase in serine phosphorylation in flagella (Koch et al. 2015), there likely exist many other ciliary PP1 targets besides PC1 and FLA8/KIF3B. Moreover, additional GSK3 targets besides PPP1R2/PP1 are presumably regulated by WNT signalling in the cilium. For example, in mouse spermatozoa, GSK3-mediated phosphorylation of septin4 is WNT-inhibited to establish a membrane barrier at the flagellar annulus (Koch et al., 2015), and septins also function in the biogenesis of primary cilia (Palander et al., 2017).

### The WNT ┫ PP1 Axis and Ciliopathy Models

The third important finding of this study is that interfering with ciliary WNT signalling recapitulates primary and motile ciliopathies: Ccny/l1 deficiency causes exencephaly, lipodystrophy, as well as male infertility in mice (this study; Koch et al., 2015). Reduced LRP6 phosphorylation and results from inhibiting WNT ligands, LRP6, and GSK3 corroborate that these ciliary defects are due to impaired WNT signalling. Rescue of the ciliary defects by inhibition of GSK3 and PP1 but not β-catenin support the involvement of WNT/GSK3 rather than WNT/β-catenin signalling. Besides ciliary WNT signalling, conventional transcriptional WNT/β-catenin signalling as well as WNT/PCP (planar cell polarity) signalling are also prominent regulators of ciliogenesis and in multiple systems, highlighting a pervasive role of this growth factor in cilia biology.

If ciliary WNT/GSK3 signalling plays a widespread role, then *LRP6* deficiency should recapitulate ciliopathies. Consistently, exencephaly and cystic kidneys are observed in *Lrp6* null mutant mice (Pinson et al., 2000) and *Lrp6* heterozygosity exacerbates pathology in a cystic kidney mouse model (Lancaster et al., 2009). LRP6 variants are associated with *spina bifida* in patients (Allache et al., 2014; Lei et al., 2015; Shi et al., 2017). Finally, mice mutant for the LRP6 kinase *Cdk16* show male infertility with sperm flagellar defects (Mikolcevic et al., 2012). Thus, we expect that the discovery how WNT ┫ PP1 signalling promotes ciliogensis will aid in the understanding of the etiology of ciliopathies.

## ACKNOWLEDGEMENTS

Expert technical help from the DKFZ core facilities for light microscopy, transgenics and the Central Animal Laboratory is gratefully acknowledged. We thank Arial Zeng for providing the Ccny mutant mice. We also thank Hyeyoon Lee for designing the graphical abstract and Sebastian Wurzbacher from the Electron Microscopy Core Facility at Heidelberg University for performing the SEM analysis. This work was supported by the Deutsche Forschungsgemeinschaft (DFG), SFB873 and SFB1324. FDS was supported by the EMBO long-term fellowship ALTF 982-2018.

## AUTHOR CONTRIBUTIONS

KZ and FDS conceived, performed and analysed mouse and human cell line experiments. JH supervised animal husbandry and assisted with experiments. MWB performed the TEM *in vivo* primary cilia analysis. WBH and CS contributed by planning experiments and revising the manuscript. CN supervised all aspects of the project. CN and FDS wrote the manuscript with input from all authors.

## DECLARATION OF INTERESTS

The authors declare no competing interests.

## MATERIALS AND METHODS

### Lead Contact and Materials Availability

Further information and requests for materials are directed to the Lead Contact, Christof Niehrs.

### Materials Availability Statement

All unique materials newly generated in this study are available from the Lead Contact.

### Experimental Model and Subject Details

#### Mice

All mouse work was conducted according to national and international guidelines and was approved by the state review board of Baden Württemberg (protocol no. G-123/18). The cyclin Y-deficient (*Ccny^-/-^*) and cyclin Y-like 1-deficient (*Ccnyl1^-/-^*) mouse lines have been described previously (Koch et al., 2015; Zeng et al., 2016). Generation of *Ccny* and *Ccnyl1* double knockout embryos (DKO) was performed by crossing *Ccny^-/-^; Ccnyl1^+/-^* males with *Ccny^+/-^; Ccnyl1^-/-^* females. *Ccny^+/-^; Ccnyl1^+/-^* embryos were used as controls. Embryos were analyzed at E13.5. For fat mass experiments, adult male *Ccny^-/-^* and *Ccny*^+/-^ control littermates were fed normal diets and sacrificed at 16 weeks. Epididymal, inguinal, and interscapular fat pads were weighed immediately after dissection. Adult mice were sacrificed by cervical dislocation.

#### Neural progenitor cell isolation and culturing

Neural progenitor cells (NPCs) were obtained by incrossing *Ccny^+/-^; Ccnyl1^+/-^* animals and collecting *Ccny^+/-^; Ccnyl1*^-/-^ (control) embryos at E13.5. The forebrain cerebral cortex was dissected from individual embryos, dissociated into a single cell suspension by repetitive pipetting, and filtered through a 70 um cell strainer. The resulting neurospheres were cultured in NPC media (DMEM/F12, B27, Glucose, Hepes, Progesterone, Putrescine, Heparin, Penicillin/Streptomycin, Insulin-Transferrin-Sodium Selenite Supplements, Sodium Bicarbonate and 20 ng/ml EGF) for 5-7 days before being passaged.

For passaging, cells were treated with accutase and then pipetted repetitively to obtain a single-cell suspension. Cells were passaged at least two times before performing experiments. For NPC monolayer cultures, single cells were grown on laminin and poly-L-lysine-coated 24 well plates for 24-72 hours before being harvested for further analysis. Knockdown of *Ccny* in *Ccny^+/-^; Ccnyl1*^-/-^ cells was performed by transducing a sh*Ccny* lentivirus (described later on) plus 4 ug/ml polybrene into neurospheres 24 hours before switching to monolayer conditions. Mature neurons were not detected in NPC cultures prior to differentiation as determined by IF for the NPC marker Pax6 (data not shown).

### Cell culture

#### Cell lines

HEK293T cells, HEK293 cells, mouse embryonic fibroblasts (MEF) and 3T3-L1 cells were cultured in DMEM supplemented with 10% v/v fetal bovine serum (fetal calf serum for 3T3-L1), 2 mM glutamine and 1% v/v penicillin/streptomycin. hTERT-RPE1 (Retinal pigment epithelial cells) were cultured in DMEM-F12 media supplemented with 10% v/v fetal bovine serum, 2 mM glutamine and 1% v/v penicillin/streptomycin. Cells were grown at 37°C and 5% CO_2_ (10% CO_2_ for 293T and 293 cells) in a humidified chamber. The *LRP5/6* DKO HEK293T cell line has been described previously (Hirsch et al., 2017).

#### Transfections

The *Xenopus* ΔN-β-catenin, human LRP6-EGFP, human LRP6*-*no tag, human ΔE1-4-LRP6, human GSK3β-Myc, human V5-FZD5, human LRP5-myc and human pArl13b-mKate2 plasmids have been previously described (Kirsch et al., 2017; Huang and Niehrs, 2014; Berger et al., 2017; Mao et al., 2001; Koch et al., 2015; Chang et al., 2020; Mao et al, 2001b; Paridaen et al., 2013) and were transfected using XtremeGENE 9 DNA Transfection reagent (Roche) at a concentration of 200 ng/ml (unless otherwise stated) according to the manufacturer’s instructions. Full length MesD (mesoderm development LRP chaperone) plasmid (Berger et al. 2017) was co-transfected with LRP5/6 plasmids at 1/10^th^ of the LRP5/6 concentration to promote LRP5/6 membrane targeting. Constitutively active *Xenopus* ΔN-β-catenin plasmid was transfected at 10 ng/ml for rescue experiments in *CCNY/L1*DKO 293T cells. For cilia rescue experiments in *LRP5/6*DKO 293T cells (Figure 3M), LRP6 (WT and CTS mutants) and LRP5-Myc constructs were transfected at 750 and 1500 ng/ml, respectively.

#### Acute siRNA/shRNA knockdowns

Scrambled and targeted siRNA SMARTpools (Horizon) and transfections were performed using Dharmafect 1 (Horizon) according to the manufacturer’s instructions. Lentiviral-mediated shRNA knockdown of *Ccny* in 3T3-L1 cells was performed as in NPCs (see above).

#### Pharmacologic and conditioned media treatment

Where indicated, cells were treated with nocodazole, Lithium Chloride (LiCl), 6-Bromoindirubin-3’-oxime (BIO), 6-[2-[[4-(2,4-dichlorophenyl)-5-(5-methyl-1Himidazol-2-yl)pyrimidin2yl]amino]ethylamino]pyridine-3-carbonitrile(CHIR99021(CHIR)) Okadaic acid (OA) and Calyculin A (Caly A) (length and dose of treatments indicated in figure legends). WNT3A and control conditioned media (CM) were prepared from stably transfected L cells (ATCC) as per the supplier’s guidelines and used at 1:5 dilution.

#### Cilia analysis

For cilia analyses in 293T, 293, MEF and RPE1 cell lines, cells were grown on poly-D-lysine coated coverslips (10 µg/ml) in 24 well plates to 60-80% confluency and then cultured in DMEM containing no serum for 24 hours (or 48 hours for 293 cells) prior to harvest. Plasmid transfections and siRNA knockdowns of targeted genes were performed one day after plating cells. For analysing 3T3-L1 cilia, cells were cultured on poly-D-lysine coated coverslips in 24 well plates at full confluency for 24 hours and then serum starved for an additional 24 hours before harvest.

#### 3T3-L1 differentiation assays

For adipocyte differentiation assays, 3T3-L1 cells were grown to confluency in DMEM containing 10% fetal calf serum, and then cultured for an extra 2 days at full confluency. Adipogenesis was induced by culturing cells for 2 days in DMEM 10% FBS media supplemented with insulin (1 mg/mL), dexamethasone (1 mM) and IBMX (0.5 mM). Media was then changed to maintenance media (DMEM containing 10% FBS and 1 mg/mL insulin) and changed every 2-3 days for a total differentiation time of 4-8 days. Where indicated, 0.5 nm Caly A was added together with the differentiation cocktail.

### Generation of *CCNY/CCNYL1* double knock-out HEK293T cells

CCNY/CCNYL1 (CCNY/L1) double knock-out HEK 293T cells were generated by CRISPR/Cas9-mediated gene editing using the following guide RNAs: *CCNY* (5’-3’GCTATCATCTAGGAAAATGG); *CCNYL1* (5’-3’ GCTGCCCTGGAGATACCTGA). The guide RNAs were chosen according to the online database E-CRISP (www.e-crisp.org/E-CRISP/). The guide sequences were cloned into a PX330-based plasmid, and were introduced into HEK293T cells by transient transfection. Following transfection, clonal colonies were obtained by limiting dilution, and deletions were confirmed by sequence analysis and immunoblot blot with anti-CCNY and CCNYL1 antibodies. Further confirmation of successful gene deletion was obtained by a WNT-reporter (SuperTOPFLASH) assay as described later on.

### Generation of *LRP5/LRP6* double knock-out MEF cells

MEF cells were isolated from E13.5 embryos carrying floxed alleles of *LRP5* and *LRP6* (Joeng et al., 2011) using standard procedures. Following isolation, *LRP5/LRP6* floxed alleles were deleted by transient transfection of a plasmid expressing Cre recombinase under the control of the CMV promoter, together with an EGFP plasmid to identify and isolate transfected colonies. Deletion of *LRP5/6* was verified by Sanger sequencing.

### Method Details Lentivirus preparation

The *Ccny* shRNA plasmid was obtained from the lab of Arial Zeng (Zeng et al., 2016) and was packaged in 293T cells according to Lois et al., 2002.

### Primary cilia isolation

Primary cilia were isolated from 293T cells using the “peel-off” technique as described in Huang et al., 2008. Briefly, 293T cells were cultured on poly-D-lysine coated coverslips until confluency. After removing media, poly-D-lysine coated cover slips were placed on the coverslips harbouring the 293T cells and then gently peeled off to remove primary cilia. Coverslips with cilia were then fixed in ice cold methanol for 10 minutes on ice and IF carried out as described later on.

### LRP6 CTS mutagenesis

Mutagenesis of the *LRP6* ciliary targeting sequences (CTSs) was performed by PCR-amplifying full length LRP6-non tagged plasmid with Phusion^TM^ DNA polymerase using primers designed to mutate selected amino acids into alanines (VP motif) or phenylalanines (AQ motif). Double VP + AQ mutations were performed sequentially.

### Transmission electron microscopy

Transmission electron microscopy (TEM) was performed as previously described (Paridaen et al., 2015). In short, mouse E13.5 embryos were fixed in 1% GA, 2% PFA in PBS, and heads or kidneys dissected and stored in 0.01% GA in PBS at 4°C until further processing.

Fixed heads were embedded in 4% low-melting agarose (ScienceServices) in 0.1 M phosphate buffer, and then sectioned at a thickness of 200 µm in the transverse orientation using a vibratome (VT1200S, Leica). Sections were post-fixed in 1% osmium tetroxide, 1.5 % potassium ferrocyanide for 30 min - 1 hour at room temperature, and subsequently contrasted with 0.5 % uranyl acetate overnight. The samples were then dehydrated through an ascending series of ethanol, and after gradual infiltration flat-embedded in Epon replacement (Carl Roth).

Fixed kidneys were processed in parallel to the head sections and split in two halves prior to resin-embedding in silicon molds. Ultrathin (70 nm) plastic sections were cut on a microtome (UCT, Leica) and post-stained with uranyl acetate and lead citrate according to standard protocols. Images were taken on a Morgagni EM at 80 kV (FEI) with a Morada camera (Olympus) and ITEM software (Olympus).

### Scanning electron microscopy

Scanning electron microscopy (SEM) was performed as previously described (Khatri et al., 2014). Briefly, whole E13.5 brains were cut in half to expose the anterior forebrain and then fixed with EM fixation buffer (4% Formaldehyde, 2% Glutaraldehyde, 10 mM Cacodylate, 1 mM CaCl_2_, 1 mM MgCl_2_, pH 7.2) for 16-24 hours at 4°C. Embryos were then washed several times and post-fixed with 2% osmium tetroxide, 1.5% potassium ferrocyanide for 1 h. Following post-fixation, samples were washed and dehydrated with an ascending series of ethanol and pure acetone before critical point drying. The samples were then sputter-coated with an 80% gold, 20% palladium alloy and examined with an ULTRA 55 field-emission scanning electron microscope (ZEISS).

### Histology and immunofluorescence

#### Paraffin sections

Tissues were fixed overnight in 4% paraformaldehyde at 4°C, progressively dehydrated and embedded in paraffin. For immunofluorescence, 7 µm thick sections were rehydrated, boiled in a pressure cooker for 2 min with Citrate/EDTA buffer (10 mM Sodium Citrate, 5 mM TrisHCL, 2 mM EDTA, pH 8.0) and blocked in PBS solution containing 5% normal donkey serum, 1% BSA and 0.1% Triton X-100. All antibodies were applied overnight at 4°C at the concentrations listed in table S1. Secondary antibodies were diluted 1:500 in PBS containing Hoechst dye (1:1000) and applied at room temperature for 1h. For Haematoxylin and Eosin staining, 7 µm thick sections from epididymal fat pads were rehydrated and stained according to standard procedures (see Da Silva et al., 2021).

#### Frozen sections

Tissues were fixed for 2 hours in 4% paraformaldehyde at 4°C, incubated in 30% sucrose overnight, embedded in Tissue-Tek (OCT, Sakura) and frozen at −20°C. 8 µm thick sections were washed briefly in PBS and then heated in a microwave for 5 minutes in Sodium Citrate buffer (10 mM Sodium Citrate, 0.05% Tween pH 6.0). Blocking and antibody applications were performed as described above.

#### Cell culture

Cells were cultured on coverslips coated with poly D-Lysine and then fixed with ice-cold methanol on ice for 10 min or 4% PFA at room temperature for 10 min. Following fixation cells were washed twice in PBS and then blocked and stained with indicated antibodies as described above. For GSK3β IF in 293T cells and endogenous LRP6 IF in MEF cells, a brief wash with cytoskeletal buffer (CB) (100 mM NaCl, 300 mM Sucrose, 3 mM MgCl_2_, 10 mM PIPES and 5 mM EGTA) was performed prior to fixation. Cells were then fixed in 4% PFA diluted in CB buffer for 10 minutes at 37°C. Subsequent blocking and antibody applications were carried out as described above. For Sp1490-LRP6 IF analysis following WNT3A treatment, 293T cells were serum starved for 48 hours and then treated with WNT3A conditioned media (diluted 1:5 in serum free media) for 30 minutes. Cells were then fixed with ice-cold methanol on ice for 10 minutes and IF carried out as described above.

For Neurosphere monolayer IF, cells were fixed in 4% PFA at 4°C for 2 hours. Following fixation, cells were washed in PBS with 0.1% Triton and blocked in blocking buffer (5% donkey serum, 3% BSA, 0.1% Tween20 in PBS) for 30 min. Subsequent antibody applications were carried out as described above.

#### Antibodies

Rabbit polyclonal antibodies against Ccny and Ccnyl1 were raised against synthetic peptides and affinity-purified as previously described (Davidson et al, 2009; Koch et al, 2015). No crossreactivity with Ccnyl1 was detected for the anti-Ccny antibody and vice versa. All other antibodies used in this study are commercial and are described in Table S1.

### Real-time quantitative PCR

mRNA from cells was extracted using the RNeasy Mini and Micro Kits (Qiagen). Extraction and precipitation of RNA were performed following the manufacturer’s instruction. Extracted mRNA was transcribed to cDNA using random hexamer primers. PCR was performed on a Roche Light Cycler 480 using the Universal ProbeLibrary system.

### Immunoblotting

Cultured cells were adjusted to equal numbers, washed with PBS, resuspended in Triton lysis buffer (20 mM Tris HCl, 150 M NaCl, 1% Triton X-100, 1 mM EDTA, 1 mM EGTA, 1 mM b-glycerolphosphate, 2.5 mM sodium pyrophosphate and 1 mM sodium orthovanadate), incubated for 30 min on ice, centrifuged at 4°C for 5 min and supernatants were collected. Lysates were heated at 70°C in NuPage LDS buffer supplemented with 50 mM DTT.

Samples were separated on polyacrylamide gels, transferred to nitrocellulose and blocked with 5% skim-milk powder or 5% BSA in Tris-buffered saline with 0.1% Tween-20 (TBST) for 1 hour at room temperature. Primary antibody was diluted in blocking buffer and incubated overnight at 4°C. Membranes were incubated with peroxidase-linked secondary antibodies for 1h at RT diluted in blocking buffer and then treated with Supersignal West Pico solution (Thermo Scientific). Images were acquired on an LAS-3000 system (Fuji Film).

### Live cell imaging

HEK293T cells were seeded at a density of 120,000 cells/well in poly-D-Lysine coated #1.5 German borosilicate Lab-Tek II 4-well chambers (Nunc) in DMEM media supplemented with 10% FBS. One day after seeding, cells were co-transfected with 200 ng/ml of pArl13b-mKate2 and LRP6-EGFP plasmids. Twenty four-hours post-transfection, medium was switched to serum-free DMEM without phenol red and cells were imaged using an LSM780 spinning disk microscope (Zeiss) equipped with an environmental chamber set to 37°C and 5% CO2. Z-stacks of 25 um were taken at 1.5 um steps, at time intervals of 15 min for a total of 24 hours.

### WNT SuperTOPFLASH assay

HEK293T cells were cultured in 96 well plates and transfected with 30 ng Super TOPFLASH (Firefly), 3 ng Renilla, 117 ng empty pCS2^+^ vector and, where indicated, 1 ng *Xenopus* ΔN-β-catenin and 5 ng LRP5/6. For testing Super TOPFLASH activation with GSK3 inhibitors, 16 hours after transfection, cells were serum starved and treated with LiCl or BIO for 24 hours. For testing Super TOPFLASH activation in CCNY/L1 single and double knockout cell lines, cells were first synchronized by overnight nocodazole treatment (100 ng/ml), and then treated with WNT3A conditioned media for 16 hours. In all experiments buffer plus vehicle was used as a control. Following treatments, cells were resuspended in passive lysis buffer and luminescence was recorded on a Fluoroskan Ascent FL luminometer (Thermo Scientific), using the Dual-Luciferase Reporter Assay system (Promega).

### Image acquisition and analysis

IF samples were imaged using a Zeiss LSM 700 microscope. The pinhole was set to 1 Airy unit at maximal optical resolution and gain and offset were calibrated so that fluorescent signal intensities were within the liner range of detection. For all analyses, ImageJ software was used to calculate various parameters such as cell number, pixel intensity and area, perimeter, and cilia length.

#### Mice

To quantify the number of primary cilia in the neocortex, immunostaining with an anti-Arl13b antibody was performed to identify the ciliary membrane, and the number of primary cilia was counted on at least five matched sections per embryo (two pictures taken per section for each brain hemisphere). Cilia number was normalized to the first three layers or cells instead of the length of the apical cortex surface to account for differences in cell density between different sections. For quantification of cilia in E9.5 ventral neural tubes, 12 µm Z-stacks at 1.5 µm steps were acquired and cilia were counted on entire Z-projections. Cilia quantification in kidney proximal tubules was performed by counting the number of cilia (Arl13b^+^) in the inner glomerular membrane (i.e. the membrane in contact with the lumen) and then normalizing to the perimeter of the luminal surface area analyzed. At least 200 cilia were analyzed per embryo. Phospho-PPP1R2^+^ cilia in the neocortex and kidney proximal tubules were identified by co-IF with an anti-Arl13b antibody and a phospho-specific (Tp72) PPP1R2 antibody. At least 200 cilia from 4-5 matched sections were counted per embryo. For all analyses, the number of embryos and litters analyzed are listed in the figure legends.

#### Cell lines

Quantification of cilia number and length in 293T, 293, RPE1 and 3T3-L1 cells was performed by IF with an anti-ARL13B antibody after 24 hours of serum starvation. Only cilia with a length greater than 0.5 µm were scored. Around 5 pictures at 40x (2x digital zoom) magnification were taken for each individual well and at least 200 cilia were counted per genotype/treatment. A similar mode was used to calculate cilia number and length in cultured NPCs, except no serum starvation was carried out. For quantifiying overepxpressed LRP6 ciliary localization, only cilia belonging to transfected cells were scored and quantifications were performed blindly. All experiments were performed in duplicates or triplicates and were repeated once or twice as indicated in the figure legends. Data are represented either as arithmetic mean of all samples for each experiment, or as arithmetic mean of a single representative experiment (indicated in figure legends). Each well was considered an independent sample.

## Data Analysis

Sample sizes (individual embryos, litter numbers, and wells (*in vitro* experiments)) are reported in each figure legend. All cell counts were performed in standardized microscopic fields using either the Fiji cell counter plug in or user-defined macros. All statistical analyses were conducted using Graphpad Prism. Data normality was tested by Shapiro-Wilk normality test and variances between groups were tested using F-test. Means between two groups were compared using two-tailed unpaired student’s *t*-test and means between multiple groups were compared using one-way or two-way analysis of variance (ANOVA) followed by Tukey’s multiple comparison tests, or with Chi square test. Statistical outliers were calculated using Grubb’s test. Results are displayed as arithmetic mean ± standard error of mean (SEM). Where indicated, results are shown as fold change vs. controls. Statistically significant data are indicated as: * p < 0.05, **p < 0.01, and ***p < 0.001. Non-significant data is indicated as ns.

---

**Figure S1, Related to Figures 1 and 2.**
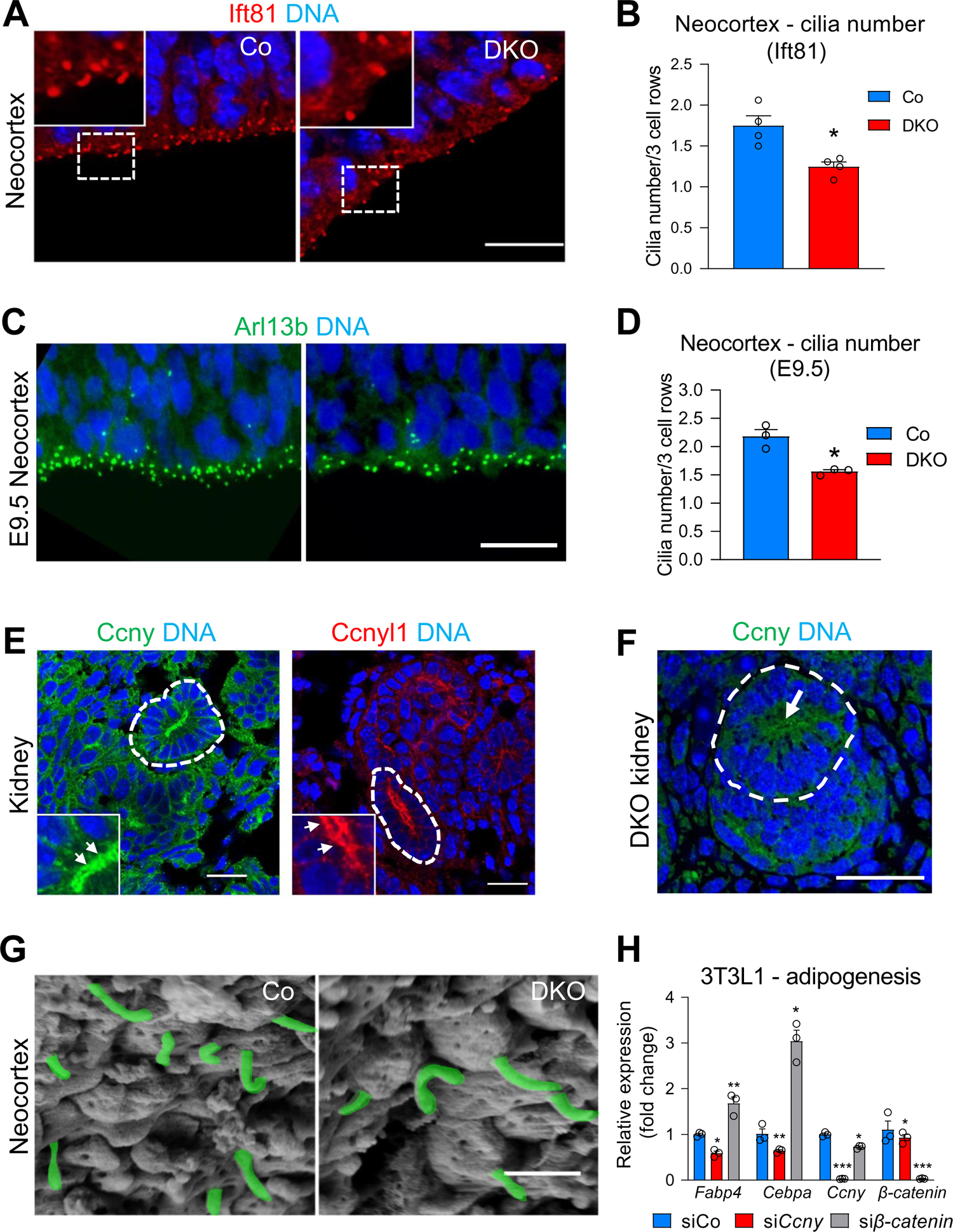
(**A-B**) **Decreased cilia number in DKO neocortex.** (**A**) IF for the ciliary axoneme protein Ift81 (Intraflagellar transport protein 81) in E13.5 neocortex of control and DKO embryos. Scale bar 10 μm. White dashed boxes are magnified in insets. (**B**) Quantification of cilia number from (**A**). Cilia quantified as in Figure 1B. Data are means ± SEM (n=4 embryos, 3 litters). (**C-D**) **Decreased cilia number in E9.5 DKO neocortex.** (**C**) IF for Arl13b in E9.5 ventral neural tube of control and DKO embryos. Scale bar 10 μm. (**D**) Quantification of cilia number from (**C**). Cilia quantified as in Figure 1B. Data are means ± SEM (n=3 embryos, 2 litters). (**E-F**) **Ccny/l1 are detected in developing kidney proximal tubules.** (**E**) Ccny and Ccnyl1 localize to developing proximal tubules (white dotted lines) in E13.5 kidneys. Insets are magnified from structures highlighted by white dotted lines. Note the apical membrane staining of Ccny/l1 (white arrows, insets). Scale bar 20 μm. (**F**) Ccny IF in DKO kidneys reveals no labeling of proximal tubule apical membrane (white arrow), other than background. (G) **Scanning Electron microscopy analysis of DKO primary cilia.** SEM analysis of the cerebral cortex in DKO forebrains reveals no major difference in the length or morphology of apical cilia. Three independent control and DKO embryos were analyzed. Cilia are colored in green for better visualization. Scale bar 2 μm. (H) **Ccny is required for adipogenesis.** qPCR analysis for the adipocyte markers *Fabp4* and *Cebp* on RNA extracted from 3T3-L1 cells treated with siRNA against *Ccny* or *β-catenin*, and then induced to differentiate into adipocytes. Data expressed as fold change vs. controls and are means ± SEM (experiment performed twice in triplicates, representative experiment shown). Data information: Unpaired two-tailed t-test for all statistical analyses: * p < 0.05, ** p < 0.01, *** p < 0.001.

**Figure S2, Related to Figure 3.**
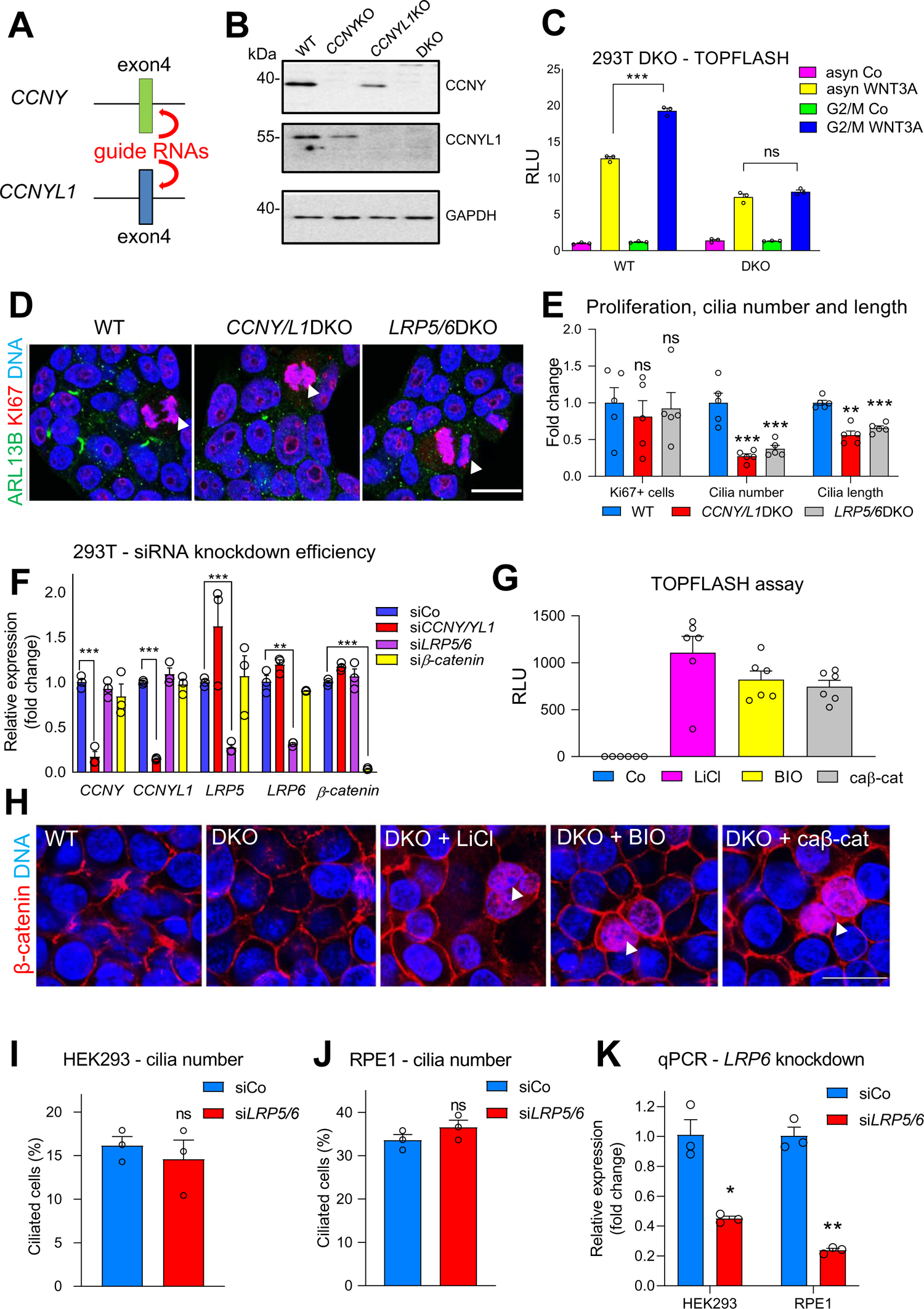
(**A-C**) **Generation and validation of 293T *CCNY/L1*DKO cells. (A)** Schematic demonstrating the strategy used to generate *CCNY* and *CCNYL1* individual knockout and *CCNY/CCNYL1* double knockout (DKO) 293T cells using CRISPR/Cas9 technology. For both genes, guide RNAs targeting exon 4 were used to generate insertion/deletion mutants. (**B**) Immunoblot with CCNY and CCNYL1-specific antibodies demonstrating the absence of CCNY/L1 proteins in single KO and DKO cells. (**C**) WNT reporter (SuperTOPFLASH) assay of single KO and DKO cells treated with 100 ng/ml nocodazole, to arrest cells in G2/M, and WNT3A-conditioned media, to activate canonical WNT signalling. Both treatments were performed simultaneously for 16 hours. Data are represented as relative luciferase units (RLU) fold change vs controls and data are means ± SEM (n=3 technical replicates). (**D-E**) **Proliferation analysis in serum-starved 293T DKO cells.** (**D**) Co-IF for ARL13B and KI67 in WT, *CCNY/L1*DKO and *LRP5/6*DKO 293T serum-starved cells. (**E**) Quantification of Ki67^+^ cells, cilia number and length from (D). Data are shown as fold change vs controls and data are means ± SEM (n=5 technical replicates). Scale bar 10μm. (**F**) **Analysis of siRNA knockdown efficiency.** qPCR analysis on RNA extracted from 293T cells treated with siRNAs against *CCNY/L1*, *LRP5/6* and *β-catenin*. Data are represented as fold change vs controls and are means ± SEM (n=3 technical replicates). (**G-H**) **Analysis of WNT signaling activation via GSK3 inhibition and *β-catenin* overexpression.** (**G)** SuperTOPFLASH assay on 293T cells treated with LiCl/BIO or transfected with constitutively active *Xenopus* ΔN-β-catenin (caβ-cat). Data are represented as RLU fold change vs. controls and are means ± SEM (n=6). (**H**) IF for β-catenin shows comparable levels of cytoplasmic and nuclear β-catenin accumulation in 293T cells treated with LiCl and BIO, or transfected with caβ-cat. Scale bar 10 μm. (**I-K**) **Ciliogenesis analysis in 293 and RPE1 cells upon si*LRP5/6* knockdown.** (**I**) Quantification of cilia number in HEK293 cells upon acute siRNA double knockdown of LRP5/6. Cells were serum starved for 48 hours for cilia analysis. Data are means ± SEM (experiment performed twice in triplicates, representative experiment shown). (**J**) Quantification of cilia number in hTERT-RPE1 (Retinal pigment epithelial) cells upon acute siRNA knockdown of LRP5/6. Cells were serum-starved for 24 hours. Data are means ± SEM (experiment performed twice in triplicates, representative experiment shown). (**K**) qPCR analysis demonstrating siRNA knockdown efficiency of *LRP6* for (I) and (J). Data are fold change vs. controls and are means ± SEM (for each cell line, experiment performed twice in triplicates, representative experiments shown). Data information: Unpaired two-tailed t-test for all statistical anaylses: ns = not significant, * p < 0.05, ** p < 0.01, *** p < 0.001.

**Figure S3, Related to Figure 4.**
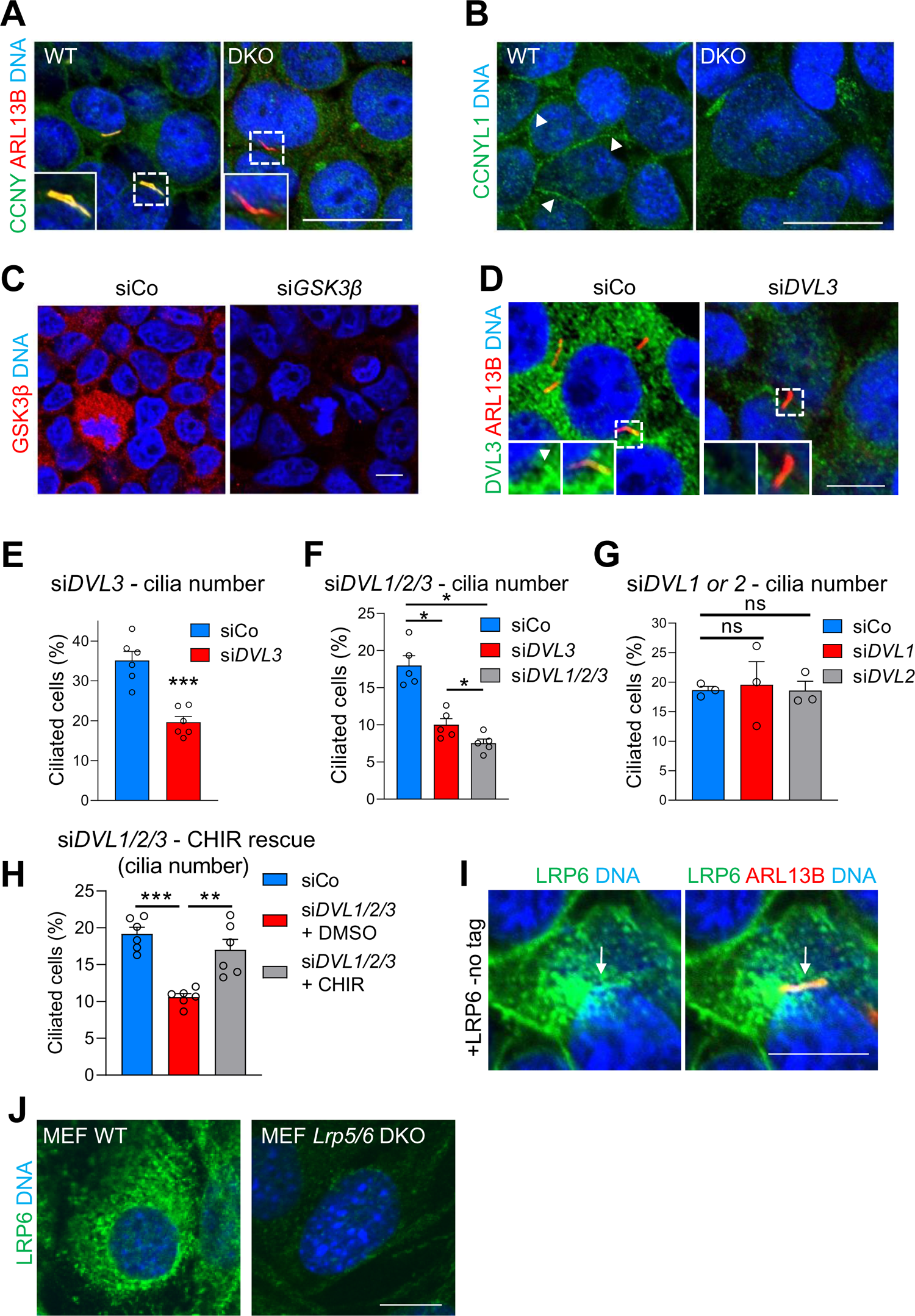
(**A-B**) **Validation of CCNY/L1 ciliary staining.** (**A**) Co-IF for CCNY and ARL13B (ARL) in WT and *CCNY/L1*DKO (DKO) 293T cells. No ciliary CCNY staining is detected in DKO cells. Areas in dashed boxes are shown at higher magnification in the insets. (**B**) CCNYL1 membrane staining (white arrowheads) is absent in DKO cells. Scale bars 10 μm. (**C-D**) **GSK3β and DVL3 ciliary staining.** (**C**) IF for GSK3β in 293T cells treated with siRNA against GSK3β to validate antibody staining. Note the strong reduction of staining in si*GSK3β*– treated cells. Scale bar 5 μm. (**D**) Co-IF for DVL3 and ARL13B in 293T cells to detect endogenous DVL3 in primary cilia. Acute siRNA knockdown of *DVL3* (right panel) performed to validate antibody staining, which, shows strong reduction in cilia (magnified in insets). Scale bars 5 μm. (**E-H**) **DVL3 is required for primary cilia formation in 293T cells.** (**E**) Quantification of cilia number in 293T cells upon siRNA knockdown of *DVL3*. Data are means ± SEM (experiment performed once in triplicate and once in duplicate, all samples shown). (**F**) Quantification of cilia number in 293T cells upon single siRNA knockdown of *DVL3* or triple knockdown of *DVL1/2/3*. Data are means ± SEM (experiment performed once in triplicate and once in duplicate, all samples shown). (**G**) Quantification of cilia number in 293T cells upon siRNA knockdown of *DVL1* or *DVL2.* Data are means ± SEM (n=3 technical replicates). (**H**) Quantification of cilia number in 293T cells upon triple siRNA knockdown of *DVL1/2/*3 followed by overnight treatment with the GSK3 inhibitor CHIR99021 (3 µM). Data are means ± SEM (experiment performed twice in triplicates, all samples shown). **(I-J) LRP6 localizes to primary cilia and validation of endogenous LRP6 staining.** (**I**) IF for LRP6 and ARL13B in 293T cells transfected with *LRP6-*no tag plasmid. White arrow depicts LRP6 localization to ciliary membrane. (**J**) Validation of endogenous LRP6 IF in WT and *LRP5/6* DKO MEF cells. Staining is absent in DKO cells. Images shown at high exposure to more clearly reveal LRP6 endogenous staining. Scale bars 5 μm. Data information: Unpaired two-tailed t-test for all statistical analyses: ns, not significant; * p < 0.05, ** p < 0.01, *** p < 0.001.

**Figure S4, Related to Figure 5.**
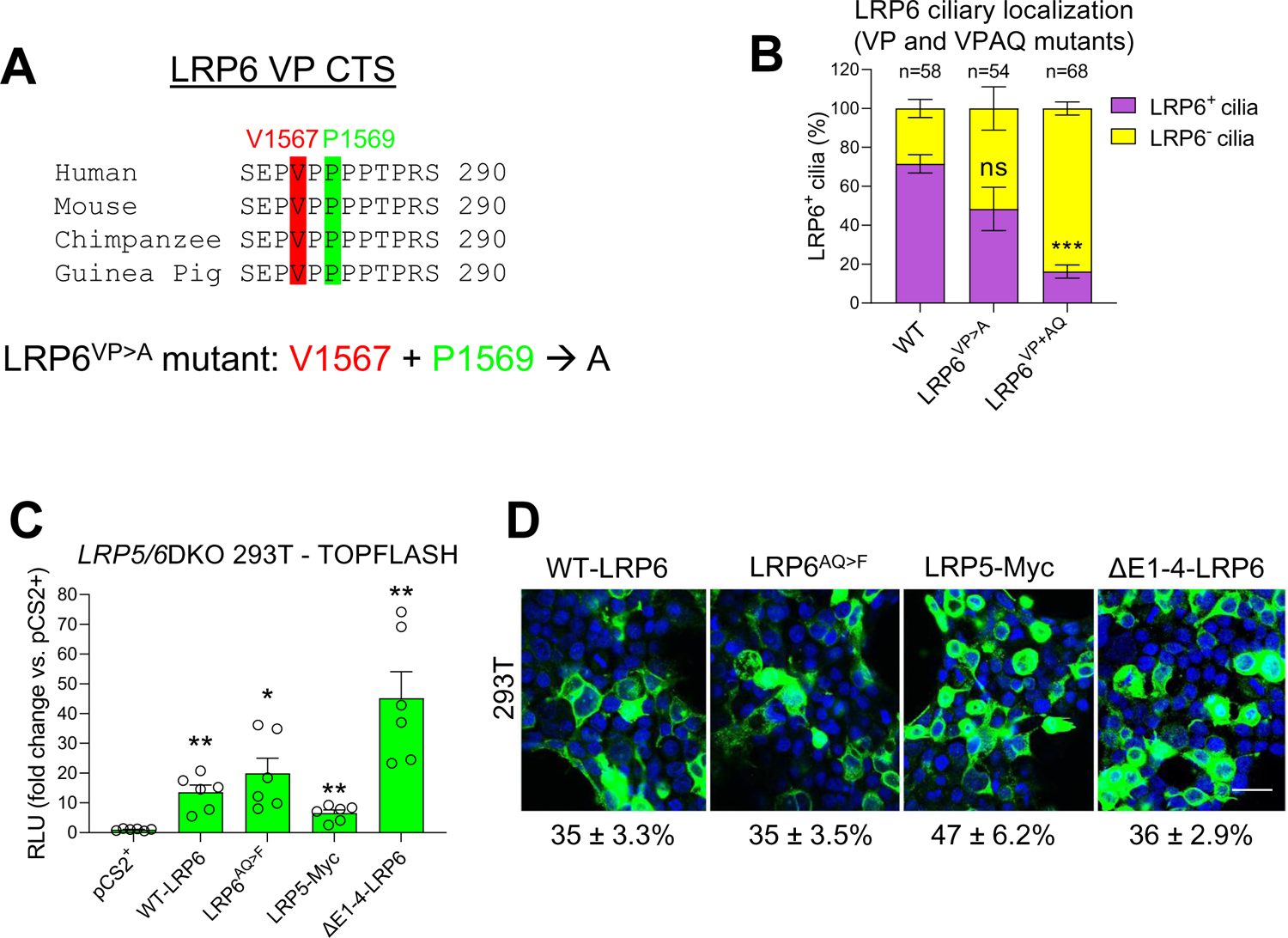
(**A-B**) **The LRP6 VP motif is dispensable for ciliary targeting of LRP6. (A)** Amino acid sequence alignment of LRP6 from human, mouse, chimapanzee and guinea pig. The VP motif is highly conserved. VP mutant form of LRP6 generated by mutating both valine (V) and proline (P) to alanine (A). (**B**) Quantification of LRP6**^+^** cilia in 293T cells transfected with WT-LRP6, LRP6^VP>A^ or LRP6^VP+AQ^ (VP and AQ mutations combined) plasmids followed by serum starvation and co-IF for LRP6 and ARL13B. Data are means ± SEM. n = number of transfected cells with visible cilia, identified by ARL13B IF. Experiment performed 3x and results pooled for final analysis. **(C-D) Ciliary localized LRP6 is required for ciliogenesis in 293T cells.** (**C**) SuperTOPFLASH assay on *LRP5/6*DKO 293T cells transfected with indicated plasmids. Data are means ± SEM normalized to pCS2^+^ (n=6 technical replicates). For statistical tests, samples compared to pCS2^+^. **(D)** IF for LRP6 (WT, AQ and ΔE1-4) or Myc (LRP5) in 293T cells transfected with indicated plasmids and serum-starved for 24 hours. Numbers represent average transfection efficiencies (normalized to cell number as determined by DNA staining) in single well (3 images per well) per plasmid. From same experiment as Figure 5F. Scale bar 15 μm. Data information: Unpaired two-tailed t-test for all statistical analyses: ns, not significant; * p < 0.05, ** p < 0.01, *** p < 0.001.

**Figure S5, Related to Figure 6.**
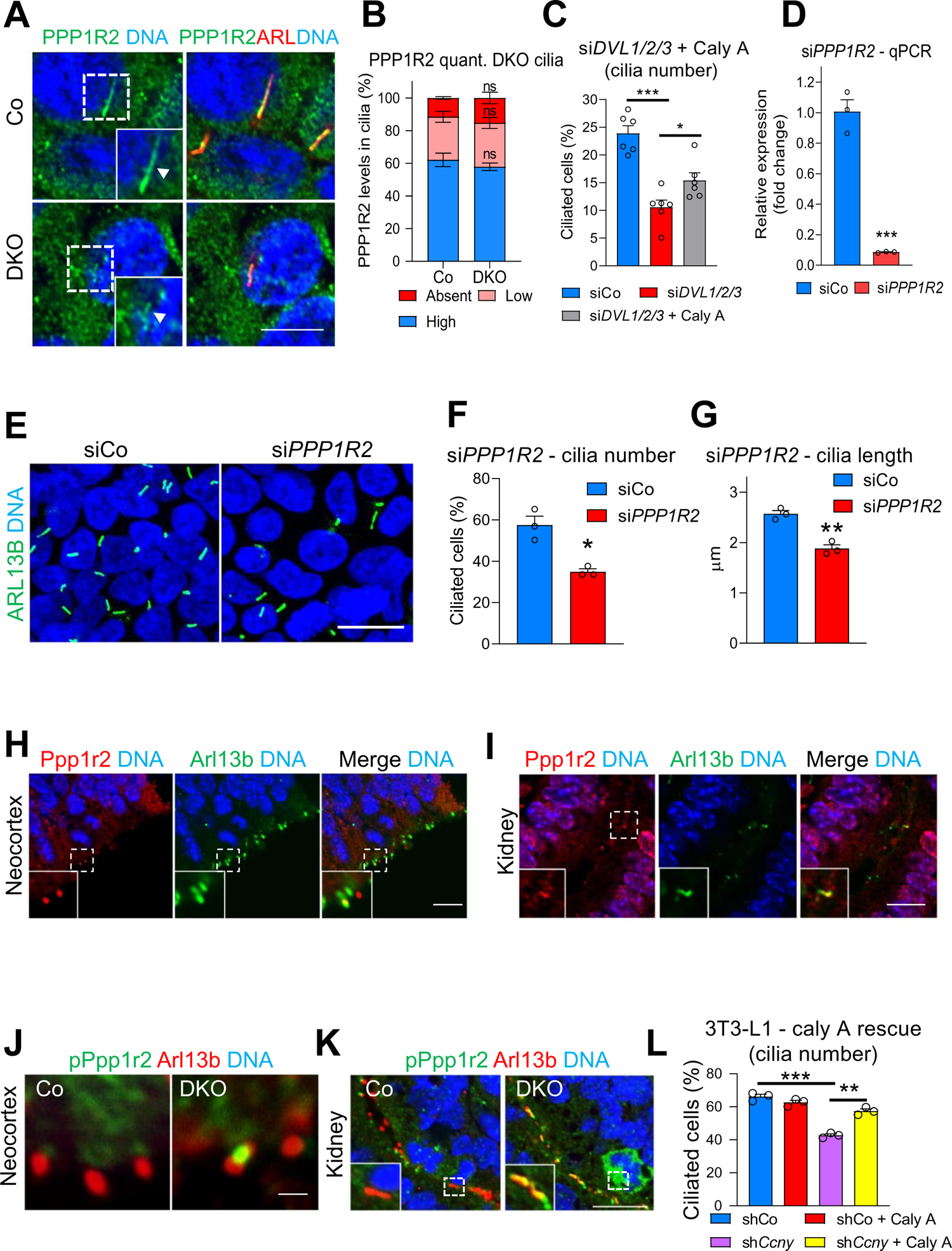
(**A-B**) **Analysis of PPP1R2 protein levels in 293T cilia**. (**A**) Co-IF for PPP1R2 and ARL13B on serum-starved 293T cells. Scale bar 10 μm. (**B**) Quantification of (A). PPP1R2 levels classified as high, low or absent based on pixel intensity. Data are means ± SEM (n=3 technical replicates). (**C**) **DVL3 regulates ciliogenesis via PP1 inhibition.** Rescue of cilia number in si*DVL1/2/3*-treated serum-starved 293T cells with the PP1 inhibitor Calyculin A (Caly A)(0.5 nM). Caly A treatment performed for 24 hours prior to harvest (i.e. during serum starvation). Data expressed as fold change vs. controls and are means ± SEM (experiment performed 2x in triplicates, all samples shown). **(D-G) PPP1R2 is required for cilia formation in 293T cells. (D)** qPCR analysis on RNA extracted from 293T cells after acute siRNA knockdown of *PPP1R2*. Data expressed as fold change vs. controls and are means ± SEM (n=3 technical replicates). (**E**) ARL13B IF in 293T cells after acute siRNA knockdown of *PPP1R2.* Scale bar 10 μm. (**F-G**) Quantification of cilia number (F) and length (G) from (G). Data are means ± SEM (for both measurements experiment performed three times in triplicates, representative experiment shown). (**H-I**) **Ppp1r2 is detected in primary cilia *in vivo*.** (**H**) Co-IF for Ppp1r2 and Arl13b in the neocortex of E13.5 embryos. Endogenous Ppp1r2 is detected in apical cilia of the neocortex (magnified in insets). Scale bar 10 μm. (**I**) Co-IF for Ppp1r2 and Arl13b in the kidneys of E13.5 embryos. Endogenous Ppp1r2 is detected in the cilia of developing proximal tubules (magnified in insets). Scale bar 10 μm. (**J-K**) **Increased phospho-Ppp1r2 levels in DKO primary cilia *in vivo*.** (**J**) Co-IF for phospho-Ppp1r2 (pPpp1r2) and Arl13b in the neocortex of E13.5 control and DKO embryos. Representative cilia shown. Scale bar 2 μm. (**K**) Co-IF for phospho-Ppp1r2 (pPpp1r2) and Arl13b in kidney proximal tubules of E13.5 control and DKO embryos. Representative cilia shown. Scale bar 10 μm. (**L**) Quantification of cilia number (identified by IF for Arl13b) in 3T3-L1 cells transduced with a lentivirus expressing sh*Ccny* and then treated with Calyculin A (0.5 nM) for 24 hours. Data are means ± SEM (experiment performed twice in triplicates, representative experiment shown). Data information: Unpaired two-tailed t-test for all statistical analyses: ns, not significant; * p < 0.05, ** p < 0.01, *** p < 0.001. Where indicated, white dashed boxes are magnified in insets.

## DOUBLE CLICK ON THE PIN TO WATCH VIDEO S1

**Video S1, related to** Figure 4: Live cell imaging of 293T cell upon exiting mitosis and forming cilia. Cells were transfected with LRP6-EGFP (green) and pArl13b-mKate2 (red) plasmids. Cells were shifted to serum-free medium at t = 0. Note co-staining of LRP6 and ARL13b (yellow) in nascent cilium (arrowhead at 3:15 h).

---

**TABLE S1:**
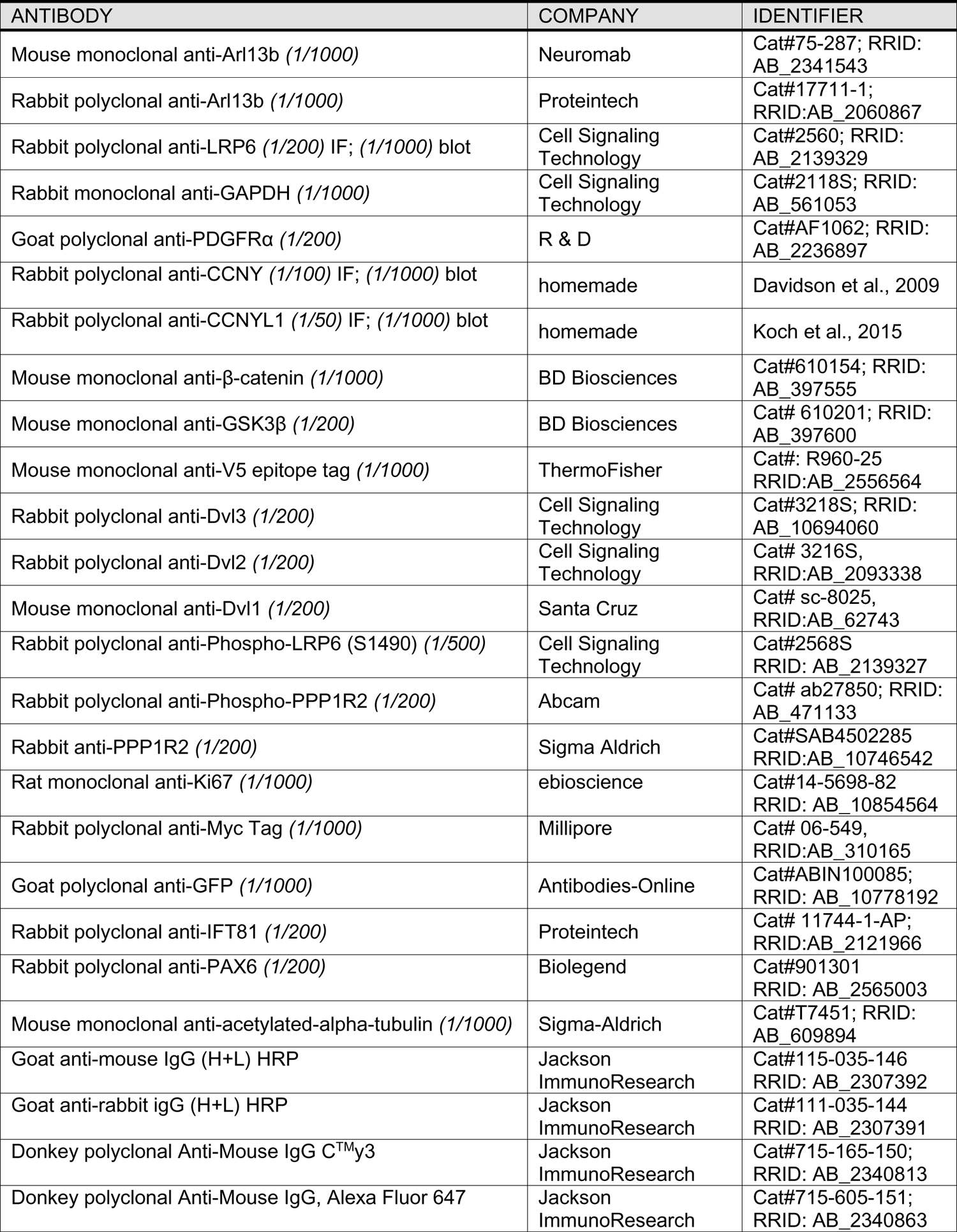
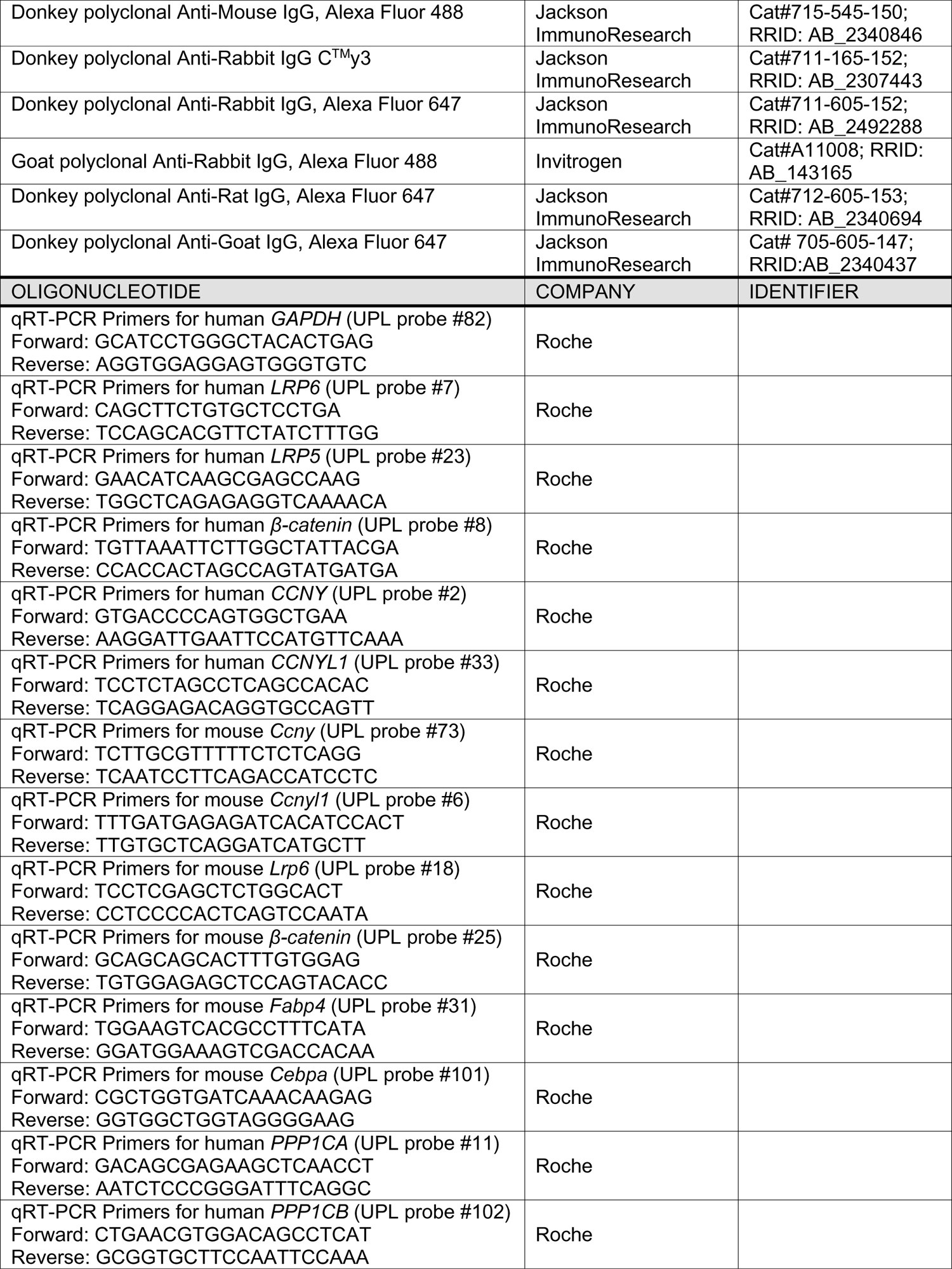
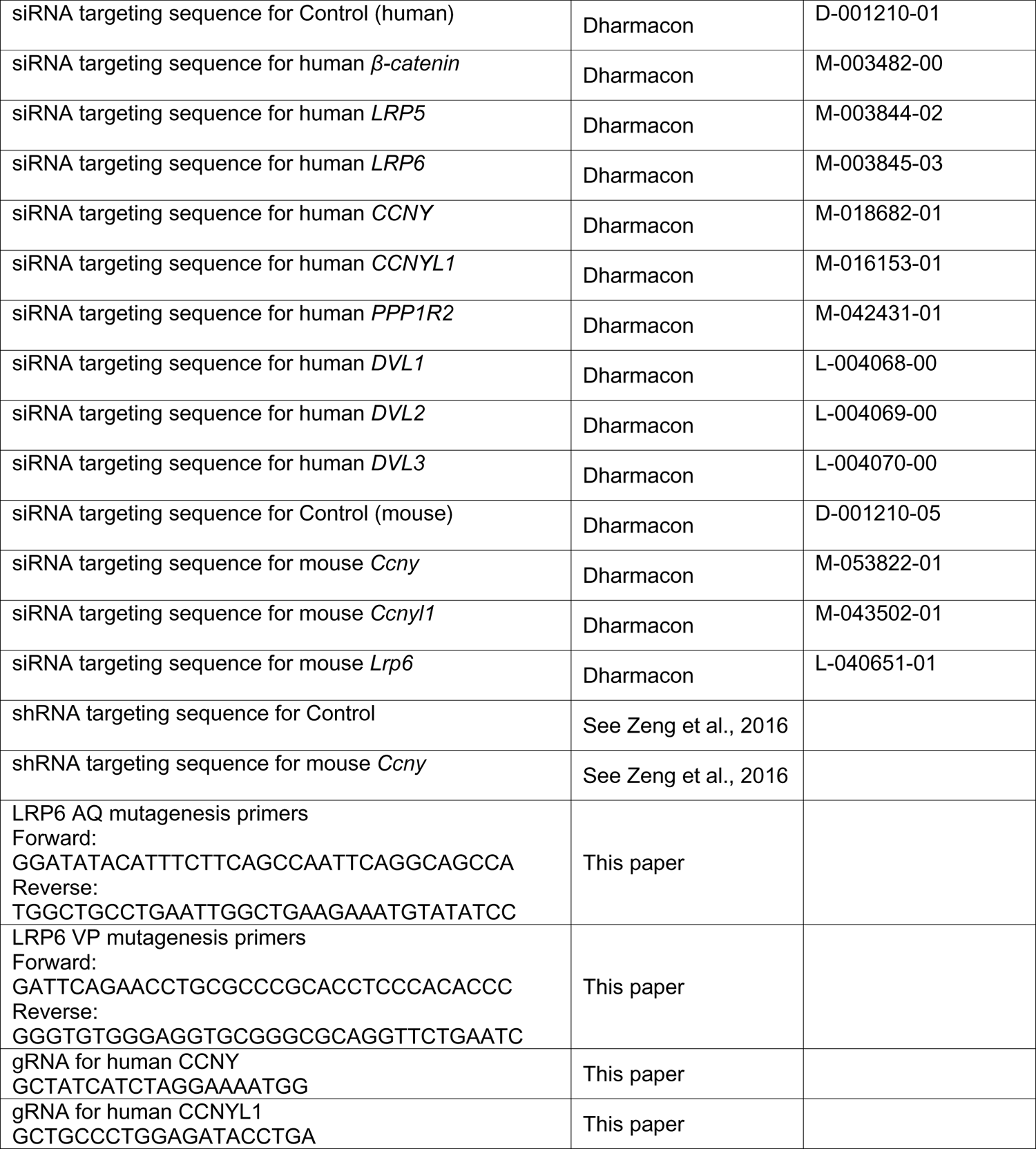
ANTIBODY AND OLIGONUCLEOTIDE TABLE

## Notes

### Competing Interest Statement

The authors have declared no competing interest.

